# HOPX governs a molecular and physiological switch between cardiomyocyte progenitor and maturation gene programs

**DOI:** 10.1101/2022.04.17.488603

**Authors:** Clayton E. Friedman, Seth W. Cheetham, Richard J. Mills, Masahito Ogawa, Meredith A. Redd, Han Sheng Chiu, Sophie Shen, Yuliangzi Sun, Dalia Mizikovsky, Romaric Bouveret, Xiaoli Chen, Holly Voges, Scott Paterson, Jessica E. De Angelis, Stacey B. Andersen, Sohye Yoon, Geoffrey J. Faulkner, Kelly A. Smith, Richard P. Harvey, Benjamin M. Hogan, Quan Nguyen, Kazu Kikuchi, James E. Hudson, Nathan J. Palpant

**Affiliations:** Institute for Molecular Bioscience, The University of Queensland, Brisbane, QLD 4072, Australia; Institute for Stem Cell and Regenerative Medicine, University of Washington, School of Medicine, Seattle, WA 98109, USA; Center for Cardiovascular Biology, University of Washington, Seattle, WA 98109, USA; Department of Medicine/Cardiology, University of Washington, Seattle, WA 98109, USA; Mater Research Institute, University of Queensland, Woolloongabba, QLD 4102, Australia; QIMR Berghofer Medical Research Institute, Brisbane, QLD 4006, Australia; School of Biomedical Sciences, The University of Queensland, St Lucia 4072, QLD, Australia; Victor Chang Cardiac Research Institute, Sydney, NSW 2010, Australia; Department of Regenerative Medicine and Tissue Engineering, National Cerebral and Cardiovascular Center Research Institute. Suita, Osaka, 564-8565, Japan; Organogenesis and Cancer Program, Peter MacCallum Cancer Centre, Melbourne, VIC 3000, Australia; Sir Peter MacCallum Department of Oncology, University of Melbourne, Melbourne, VIC 3000, Australia; Genome Innovation Hub, The University of Queensland, Brisbane, QLD 4072, Australia; Queensland Brain Institute, University of Queensland, Brisbane QLD 4072, Australia; Department of Anatomy & Physiology, The University of Melbourne, Parkville, VIC, 3010, Australia; St. Vincent’s Medical School and School of Biotechnology and Biomolecular Science, UNSW Sydney, NSW 2054, Australia; Organogenesis and Cancer Program, Peter MacCallum Cancer Centre, Melbourne, VIC 3000, Australia; Sir Peter MacCallum Department of Oncology, University of Melbourne, Melbourne, VIC 3000, Australia; Department of Anatomy and Physiology, University of Melbourne, Melbourne, VIC 3010, Australia

**Keywords:** cardiomyocyte differentiation, human pluripotent stem cell, HOPX, CAGE-seq, DamID-seq, heart development, cell proliferation, CRISPRi

## Abstract

This study establishes the homeodomain only protein, HOPX, as a determinant controlling the molecular switch between cardiomyocyte progenitor and maturation gene programs. Time-course single-cell gene expression with genome-wide footprinting reveal that HOPX interacts with and controls core cardiac networks by regulating the activity of mutually exclusive developmental gene programs. Upstream hypertrophy and proliferation pathways compete to regulate *HOPX* transcription. Mitogenic signals override hypertrophic growth signals to suppress *HOPX* and maintain cardiomyocyte progenitor gene programs. Physiological studies show HOPX directly governs genetic control of cardiomyocyte cell stress responses, electro-mechanical coupling, proliferation, and contractility. We use human genome-wide association studies (GWAS) to show that genetic variation in the HOPX-regulome is significantly associated with complex traits affecting cardiac structure and function. Collectively, this study provides a mechanistic link situating HOPX between competing upstream pathways where HOPX acts as a molecular switch controlling gene regulatory programs underpinning metabolic, signaling, and functional maturation of cardiomyocytes.

## INTRODUCTION

During heart development, molecular and physiological cues underlying organ morphogenesis guide cardiomyocyte differentiation to increase organ size and performance through carefully orchestrated gene programs governing cell maturation(Alexanian et al., 2017; Eulalio et al., 2012; Palpant et al., 2013; Ueno et al., 2007). Cardiomyocyte maturation involves regulation of cell cycle activity, changes in metabolic energy use, and significant alterations in cell structure and function(Karbassi et al., 2020). Cardiac gene programs underlying these changes are mediated by complex chromatin-level changes(Sim et al., 2021) coupled with heterotypic transcription factor interactions involving DNA regulatory proteins including GATA factors, T-box factors, MADS box transcription enhancer factors, and homeobox transcription factors(Luna-Zurita et al., 2016).

The homeodomain only protein X, HOPX, is a non-DNA binding homeodomain protein that likely acts as a regulatory scaffold protein that interacts with cardiac-associated transcriptional and chromatin factors including SRF, GATA factors, SMADs, and HDACs to guide gene programs during heart development(Trivedi et al., 2010). *Hopx* null mice succumb to partially penetrant lethality(Chen et al., 2002; Shin et al., 2002) and those surviving to adulthood have enlarged hearts due to cardiomyocyte hyperplasia(Shin et al., 2002). Reciprocally, forced HOPX expression results in cardiac hypertrophy(Friedman et al., 2018; Kook et al., 2003) and dysregulation of HOPX is associated with heart failure(Trivedi et al., 2011).

Beyond cardiac development, HOPX plays a critical role in development and homeostasis of diverse tissues(Berg et al., 2019; Loh et al., 2016; Palpant et al., 2017a) and is implicated in diseases including cancer where it has dual roles as both an oncogene and tumor suppressor in different malignancies(Chen et al., 2015; Lin et al., 2020; Pavlova et al., 2021). Evidence to date suggests the diverse functions of HOPX in development and disease are not only related to its promiscuity with diverse interaction partners but also linked to selective activation of *HOPX* alternative promoters(Friedman et al., 2018; Pavlova et al., 2021). HOPX is a critical molecule with complex roles in development and disease. However, little is known about transcriptional control of *HOPX* and how *HOPX* regulated gene programs influence changes in cardiomyocyte cell identity and function during differentiation.

This study uses single cell genomics, genome-wide footprinting, and human genetics linked with *in vitro* and *in vivo* genetic models of HOPX gain and loss of function to situate HOPX as a bridge between two mutually exclusive cardiac gene programs governing heart development. We identify upstream and downstream mechanisms controlling *HOPX* transcription and its regulatory network as a molecular switch governing functional, metabolic, and structural phenotypes of cardiomyocytes at the cell, tissue, and organ level. We use complex trait genetic data to link *HOPX* regulatory networks as dominant drivers of the genetic variation underpinning population-level changes in cardiac structure and function. Collectively, these new mechanistic insights reveal the molecular basis of *HOPX* control of cardiomyocyte identity. The work contributes to understanding fundamental biology of the heart and advances our knowledge of molecular targets governing heart disease and regeneration, processes determined by molecular control of cell growth and proliferation.

## RESULTS

### *Hopx* is associated with gene programs governing cardiomyocyte maturation

To deconstruct cell-specific regulatory networks governing cardiomyocyte differentiation during heart development *in vivo*, we used label transferring to integrate time course single cell data of mouse development between embryonic day 6.5 to postnatal day 21 (E6.5-P21)(DeLaughter et al., 2016; Lescroart et al., 2018; Li et al., 2016) (**Figure 1A-B**). We evaluated temporal-genetic changes associated with transitions during heart development across 2,337 annotated cardiomyocytes (**Figure 1B-D**). As expected, genes involved in myofibrillar maturation (*Myl2, Myh6*) and the fetal cardiac gene program (*Nppb*) increase during development while genetic markers of cardiomyocyte proliferation (*Cdk4, Ccnb1*) and mevalonate pathway activity (*Sqle, Hmgcs1*) decrease during development (**Figure 1D**). *Hopx* is detected in 89% of cardiomyocytes beginning in E8.5 cardiac progenitor cells consistent with its known role in early stages of cardiac specification(Jain et al., 2015) (**Figure 1C-D**). During cardiac chamber formation and septation (E10.5-11.5), *Hopx* expression increases to a steady state during the mid-gestation to postnatal stages of development.

**Figure 1.**
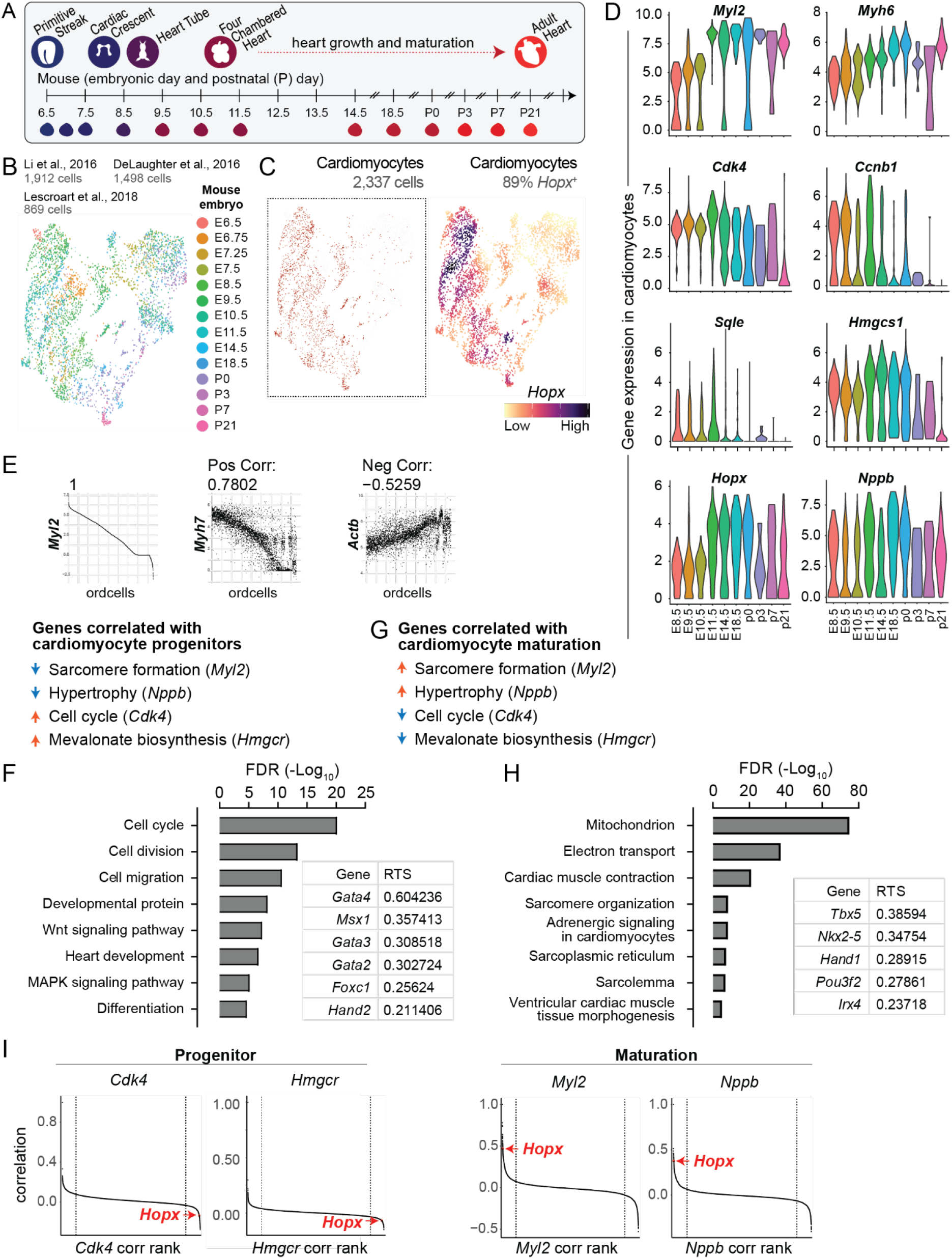
*Hopx* expression correlates with gene programs governing cardiomyocyte maturation *in vivo*. **(A)** Schematic of heart development stages represented in single cell RNA-seq data sets during a time course of mouse heart development from embryonic day (E) 6.5 to postnatal day (P) 21. **(B)** UMAP of mouse embryonic heart development single cell RNA-seq data integrated into a single gene expression data matrix. (**C-D**) UMAP analysis of *Hopx*-expressing cardiomyocytes across the developmental time course (**C**) and violin plots of cardiomyocyte-specific gene expression over time for *Hopx* and genes involved in cardiac myofibrillogenesis (*Myl2, Myh6*), cell cycle regulation (*Cdk4, Ccnb1*), mevalonate pathway (*Sqle, Hmgcs1*), and cell growth (*Nppb*) (**D**). (**E-F**) Analysis of 2,337 annotated cardiomyocytes between E6.5-P21 reveals genes negatively correlated with expression of *Myl2* and *Nppb* but positively correlated with expression of *Cdk4* and *Hmgcr* (**E**). Example correlation analysis: genes ranked by abundance for *Myl2* gene expression (left) compared with transcript abundance of *Myh7* and *Actb* using exact cell rank order for *Myl2*. Across 2,337 cardiomyocytes, *Myh7 and Actb* expression is strongly correlated with *Myl2*. (**F**) 4,225 cardiac progenitor-associated genes identified by the selection criteria in (**E**) are enriched in GO biological processes governing early heart development. Inset ranks genes with high repressive tendency scores (RTS) (Shim et al., 2020) suggesting enrichment of known developmental cardiac regulatory genes in the gene set. (**G-I**) Analysis of 2,337 annotated cardiomyocytes between E6.5-P21 reveals genes positively correlated with expression levels of *Myl2* and *Nppb* but negatively correlated with expression levels of *Cdk4* and *Hmgcr*. (**H**) 1,428 cardiac maturation genes are enriched in biological processes associated with heart maturation and function. Inset ranks genes with high repressive tendency scores (RTS)(Shim et al., 2020) suggesting enrichment of known ventricular maturation regulatory genes. (**I**) Gene rank correlation data for progenitor markers *Cdk4* and *Hmgcr* and maturation markers *Myl2* and *Nppb* in 2,337 annotated cardiomyocytes between E6.5-P21 identifies *Hopx* as negative correlated with progenitor gene programs and positive correlated with maturation gene programs.

We evaluated temporal dynamics of gene expression in thousands of cardiomyocytes across this time course of heart development between E8.5-P21 to identify positive gene sets distinguishing cardiac progenitor and maturation gene programs. The cardiac progenitor gene program was determined by identifying genes positively correlated with cell cycle (*Cdk4*) and mevalonate pathway (*Hmgcr*) and negatively correlated with cell growth (*Nppb*) and myofibrillar maturation (*Myl2*). We identified 4,225 shared genes (**Figure 1E, Figure S1A, Table S1**) enriched in gene ontologies associated with cardiac progenitor biology including Wnt and MAPK signaling, heart development, and proliferation (**Figure 1E-F**). We used a method that predicts cell regulatory genes from any orthogonal gene expression data set from any cell or tissue type (TRIAGE (Shim et al., 2020)) to identify the dominant molecular regulators of these gene programs. The top ranked regulatory genes in cardiac progenitor cells include known regulators of mesoderm specification such as GATA factors, *Msx1*, and the second heart field marker, *Hand2* (**Figure 1F**).

Genes associated with cardiomyocyte maturation were identified using the same approach but with reciprocal correlations with reference genes; namely, genes positively correlated with myofibrillar maturation (*Myl2*) and cell growth (*Nppb*) and anticorrelated with cell proliferation (*Cdk4*) and mevalonate pathway activity (*Hmgcr*) (**Figure 1G, Figure S1B, Table S1**). These data revealed 1,428 genes representing a positive gene set for cardiac maturation that are enriched in biological processes critical for cardiac cell differentiation including mitochondrial biogenesis, muscle contraction, and adrenergic signaling (**Figure 1G-H**). Expected cardiac maturation transcription factors identified by TRIAGE including *Tbx5, Nkx2-5*, and *Irx4* were among the top ranked regulatory genes (**Figure 1H**). These data establish cardiac progenitor and maturation gene programs with anticorrelated expression dynamics during mouse heart development and reflect distinct regulatory processes governing cardiac cell differentiation.

We evaluated *Hopx* expression by reference to its correlation with *Cdk4, Hmgcr, Myl2*, and *Nppb* (**Figure 1I**). We found *Hopx* within the top 10% of genes negatively correlated with *Cdk4* and *Hmgcr* while it was among the most positively correlated with *Myl2* and *Nppb*. This indicates that *Hopx* transcriptional abundance is associated with down-regulation of progenitor gene programs and up-regulation of cardiomyocyte maturation gene programs during heart development (**Figure 1I**).

### HOPX interactions across the genome bridge cardiac progenitor and maturation gene programs

While Hopx is a non-DNA binding homeodomain protein, it interacts with diverse transcriptional regulators(Jain et al., 2015; Trivedi et al., 2010) to orchestrate cardiac gene programs. We therefore set out to identify the landscape of HOPX genetic interactions in human pluripotent stem cell (hiPSC) derived cardiomyocytes. Human iPSCs provide a scalable strategy for differentiation of cardiomyocytes and recapitulate gene programs governing *in vivo* heart development(Friedman et al., 2018) (**Figure 2A-B**). We used DNA adenine methyltransferase identification (DamID) sequencing(Aughey et al., 2019; Cheetham et al., 2018; van Steensel and Henikoff, 2000) to detect HOPX occupancy genome-wide in hiPSC-derived cardiomyocytes (**Figure 2C**).

**Figure 2.**
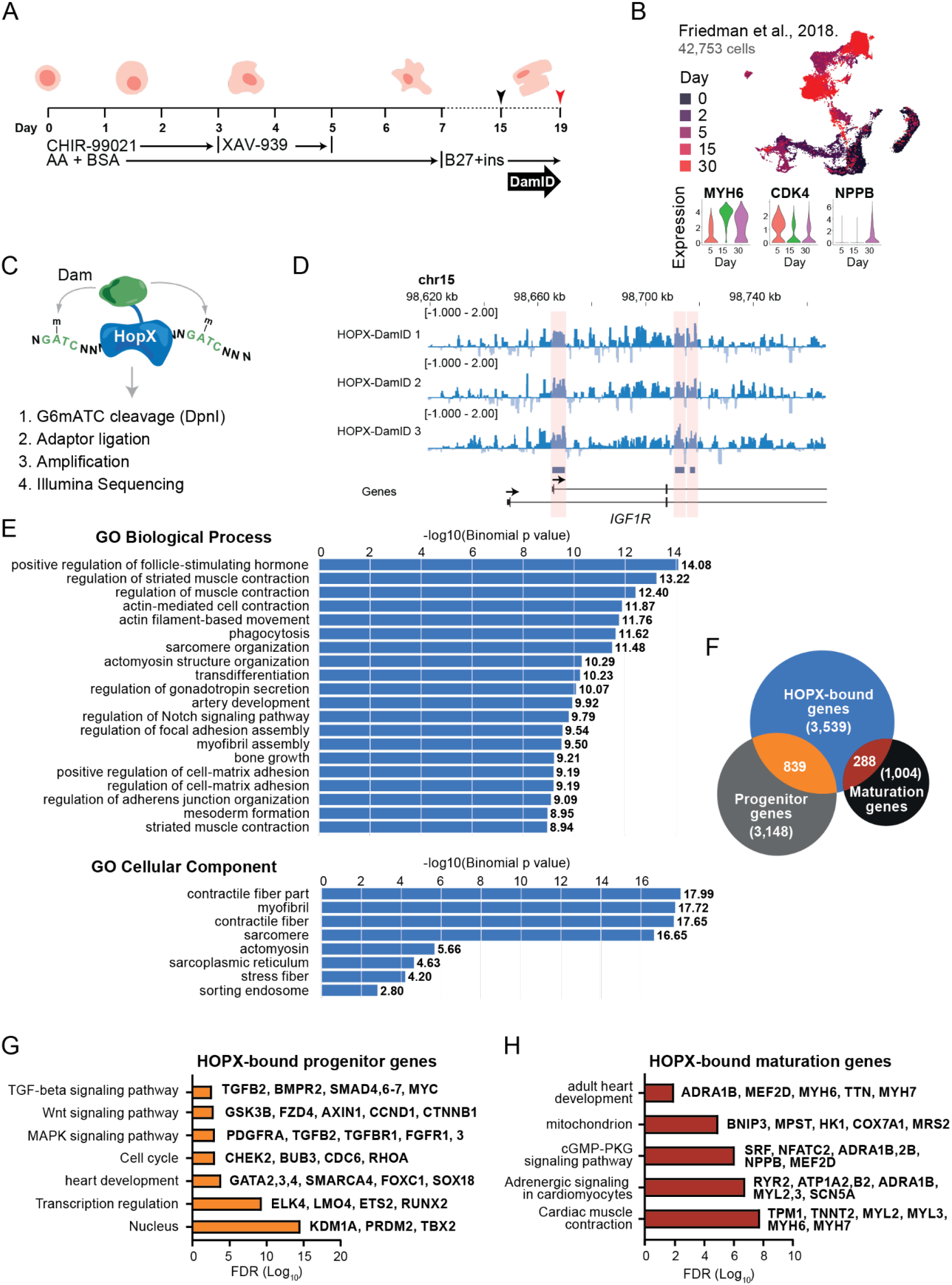
Footprinting by DamID-sequencing reveals a central role for HOPX in regulating cardiomyocyte developmental gene programs. (**A-C**) hiPSC differentiation into cardiomyocytes using small molecule Wnt modulation (**A**) recapitulates genetic programs of heart development *in vivo* as represented by time course single cell analysis (**B**). Black arrow demarcates day 15 replating into low density monolayer cultures to induce cell growth conditions with lentiviral-mediated HOPX footprinting by DNA adenine methyltransferase identification (DamID) sequencing (**C**) measured on day 19 of differentiation. **(D)** Representative HOPX footprint DamID-seq replicates and consensus peaks (peaks present in all replicates) at the *IGF1R* locus. Genome-wide DamID-seq analysis in iPSC-derived cardiomyocytes was derived from *n* = 3 replicates as shown in representative data. **(F)** Gene ontology analysis of biological processes and cellular components associated with HOPX-bound sites reveal enrichment of cardiac differentiation and function. (**F-H**) Venn diagram analysis of HOPX-bound loci in cardiomyocytes versus *in vivo* selected cardiac development gene programs identified in **Figure 1E-H** (**F**) reveals HOPX interacts with genetic loci related to cardiac progenitor (**G**) and maturation (**H**) gene programs.

We generated lentiviruses in which the Dam adenine methyltransferase gene was fused in-frame after the consensus *HOPX* coding DNA sequence. Human iPSC-derived cardiomyocytes were transduced with Dam-alone or HOPX-Dam lentiviruses in triplicate. Genomic DNA, including adenine-methylated DNA within GATC palindromic sequences, was isolated after 72 hours for DamID-seq. Next generation sequencing detected 2,784 binding sites present in all replicates (**Figure S2, Table S2**) with **Figure 2D** showing example HOPX binding sites at the *IGF1R* locus, a key regulator of cardiomyocyte hypertrophy(Ock et al., 2016). Gene ontology analysis of HOPX-bound loci revealed significant enrichment of genes associated with muscle contraction and mesoderm formation, consistent with the known role of HOPX in cardiomyocyte differentiation(Friedman et al., 2018) (**Figure 2E, Table S2**).

By reference to gene sets underpinning *in vivo* cardiomyocyte differentiation gene programs (**Figure 1E-H, Figure S1A-B**), DamID-seq data show that HOPX interacts with genes associated with both progenitor and maturation networks (**Figure 2F**). HOPX bound to 839 cardiac progenitor-associated genes which were enriched in gene ontologies associated with heart development transcription regulation (*GATA2-4, ELK4*) as well as central regulators of Wnt (*CTNNB1*), MAPK (*TGFB2*), and TGFβ (*MYC*) signaling pathways (**Figure 2G, Table S2**). HOPX also bound to 288 cardiac maturation-associated genes which enriched in biological processes including mitochondrial biogenesis (*BNIP3*), contractility (*MYL2*), adrenergic signaling (*ADRA1B*), and adult heart development (*MEF2D, NPPB*) (**Figure 2H, Table S2**). Taken together, these data demonstrate that HOPX physically interacts across anticorrelated cardiac progenitor and maturation genetic programs and suggests that HOPX operates as a central mediator of this cell state transition.

### HOPX functionally regulates cardiac progenitor and maturation gene programs

We used genetic loss-of-function studies to evaluate the role of HOPX in governing cardiac progenitor and maturation gene programs. CRISPRi iPSCs(Mandegar et al., 2016) were genetically engineered with gRNAs targeting the distal *HOPX* transcriptional start site (Friedman et al., 2018) to induce transcriptional repression of *HOPX* in a doxycycline (DOX) dependent manner. *HOPX* CRISPRi cells were differentiated into cardiomyocytes ± DOX and analyzed by genome-wide cap analysis gene expression (CAGE) sequencing (**Figure 3A-B, Figure S3, Table S3**). Representative raw DamID-seq data and CAGE-seq peaks are shown for the *MYH7* locus (**Figure 3C**). We identified 255 HOPX-interacting loci (DamID-seq) that are repressed in a HOPX-dependent manner (**Figure 3D-F, right, Table S3**). These genes are enriched in factors associated with cardiac progenitor cell biology including *CTNNB1, MYH6, SRF*, and *KDR*.

**Figure 3.**
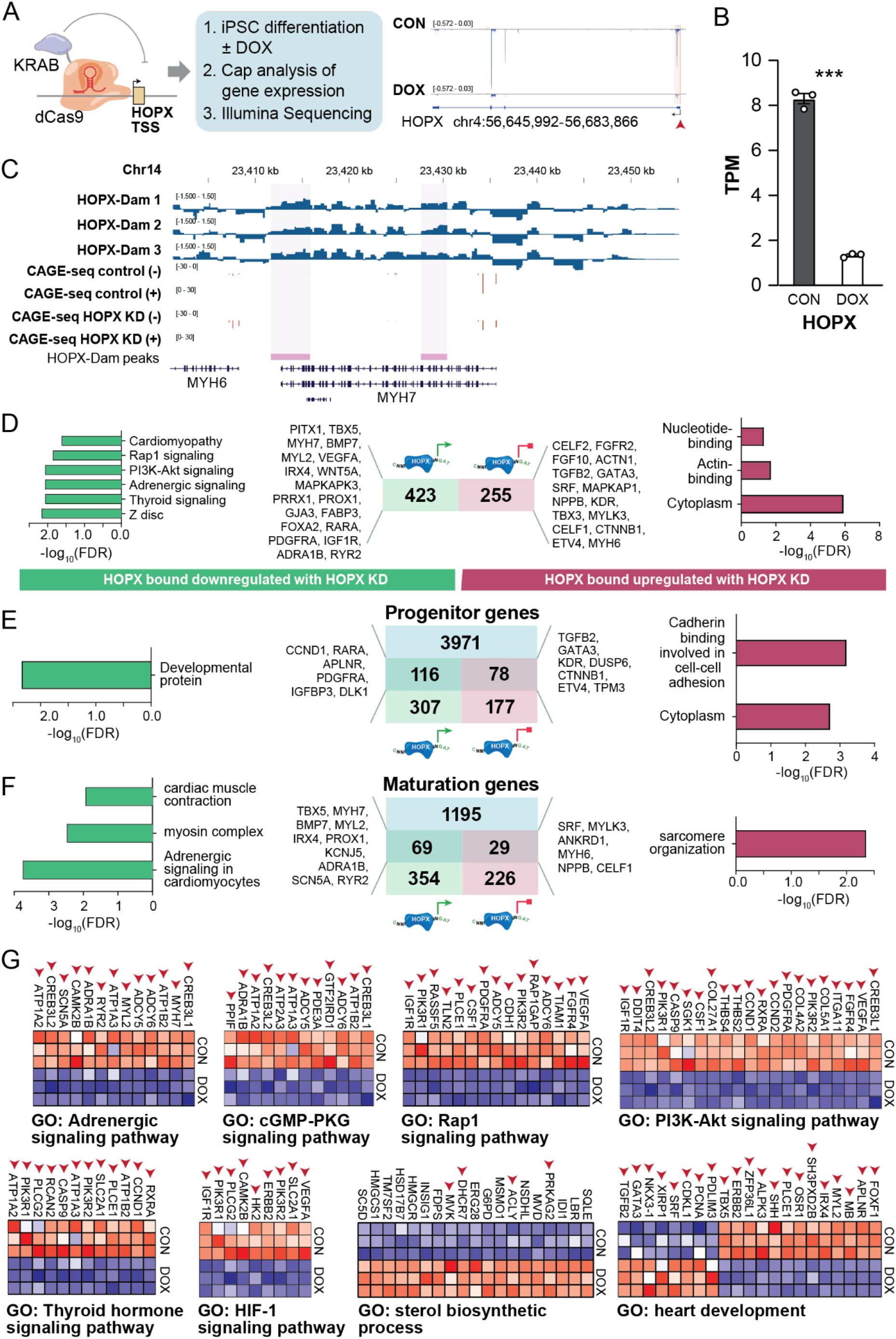
Genetic loss-of-function reveals *HOPX*-dependent cardiomyocyte maturation. (**A-B**) CRISPRi *HOPX* loss-of-function hiPSC (**A**) results in significant reduction in *HOPX* transcription in differentiated cardiomyocytes (**B**). Representative data showing DamID-seq alignment with CAGE-seq peaks demonstrating robust *HOPX* knockdown after doxycycline treatment (DOX) relative to control (CON). Statistical significance tested using student’s t-test with Welch’s correction. Error bars show mean ± S.E.M. P<0.001 (***). CAGE-seq analysis in hiPSC-derived cardiomyocytes was performed across *n* = 3 biological replicates. **(C)** Representative DamID-seq data combined with merged CAGE-seq analysis of + and - strand reads at the *MYH7* locus derived from control or *HOPX* knockdown (KD) cardiomyocytes. **(D)** Gene ontology analysis of HOPX-bound genes upregulated (right) or downregulated (left) with *HOPX* loss-of-function in differentiated cardiomyocytes reflecting genes that are transcriptionally repressed by HOPX (right) or transcriptionally dependent on HOPX (left). (**E-F**) HOPX-bound loci that are up- or downregulated and overlap with gene programs associated with *in vivo* cardiac progenitor (**E**) or maturation (**F**) gene programs as identified in **Figure 1E-H.** (**G**) Heat maps showing CAGE-seq abundance of genes in control (CON) versus *HOPX* KD (DOX) among gene ontologies associated with functional and signaling control of cardiac development. Arrows demarcate HOPX-bound genes based on DamID-seq. Heatmaps display z-scores [(sample TPM – average TPM)/standard deviation of average TPM] calculated on a gene-by-gene basis for genes enriched under annotated GO terms.

We also identified 423 HOPX-bound loci that are activated in a HOPX dependent manner (**Figure 3D-F, left, Table S3**). HOPX-dependent loci are enriched in critical determinants of heart development including transcription factors (*TBX5, IRX4*) as well as myofibrillar isoforms associated with maturation (*MYH7, MYL2*), genes involved in electromechanical coupling and chronotropic control of cardiac physiological performance (*RYR2, ADRA1B, KCNJ5, SCN5A*), and signaling pathways underpinning metabolic and signaling control of heart development (*VEGFA, PDGFRA, RXRA, ERBB1-2, IGF1R, PLCE1, SLC2A1, PIK3R1*) (**Figure 3D-F, left, Table S3**).

Based on this, we analyzed CAGE-seq data for broader changes in gene programs differentially expressed between *HOPX* KD (DOX) and control (CON) cardiomyocytes. While HOPX binds at genomic loci to regulate a subset of cardiac target genes (**Figure 2F**), CAGE-seq identified the entire cardiac progenitor- and maturation-associated repertoire of biological processes regulated in a HOPX-dependent manner (**Figure 3G**). Specifically, *HOPX* KD resulted in down-regulation of key development signaling pathways including thyroid hormone, adrenergic, cGMP, HIF-1, PI3K-Akt, and RAP1 pathways. Genes involved in the progenitor-associated sterol biosynthetic pathway, for which HOPX only binds at 4 pathway-associated loci, was broadly up-regulated with *HOPX* KD (**Figure 3G, Table S3**). Key factors identified with cardiac development similarly showed a binary on/off switch in a HOPX-dependent manner.

To assess putative HOPX-associated factors regulating these biological processes, we performed motif analysis of DamID-seq peaks using MEME-ChIP(Machanick and Bailey, 2011). DamID-seq peaks at genes activated by HOPX (downregulated in *HOPX* KD) and repressed by HOPX (upregulated in *HOPX* KD) were separated into groups. *HOPX*-repressed genes were enriched in motifs for known cardiac pacemaker transcription factors (*TBX3*) and Hippo signaling (*TEAD4*), while *HOPX*-activated genes were enriched in motifs for steroid hormone receptors (*ANDR, PATZ1*) and cardiac-associated transcription factors (*FOXJ3*) (**Figure S4A-C**).

Collectively, time-course data on heart development identified cardiac progenitor and maturation gene programs that are transcriptionally anticorrelated. These gene programs encode biological processes that appear genetically mutually exclusive, potentially controlled as binary switches during cardiac cell differentiation. We show that HOPX is a central switch mechanism governing signaling pathways, metabolic programs, regulatory factors, and structural gene programs bridging the transition between progenitor and maturation states during cardiac cell differentiation.

### Cell hypertrophy and mitogenic signals compete for regulation of *HOPX* transcription

We hypothesized if HOPX regulates a switch mechanism governing cardiac progenitor versus maturation gene programs, then transcriptional control at the *HOPX* locus must rely on competitive signaling cues that operate between these two states. *HOPX* has two transcriptional start sites with the distal TSS showing dominant control of the *HOPX* locus in cardiomyocytes(Friedman et al., 2018). These promoters show tissue-dependent epigenetic features including an active promoter region, strong transcription, as well as enhancers overlapping sites of open chromatin (ATAC-seq) across the full time course of heart development(Gorkin et al., 2020) (**Figure S5**).

Analysis of the human *HOPX* genomic locus shows diverse isoforms with GeneHancer revealing interactions between cis regulatory elements and intragenic regions (**Figure 4A-B**). GTEx data reveal cardiac specific *HOPX* isoforms (**Figure S6**) and expression quantitative trait loci (eQTL) variants spanning the *HOPX* locus in diverse tissues. Left ventricular-specific eQTLs are situated exclusively within 10kb of the distal *HOPX* TSS, overlapping motifs of key regulators of cardiac maturation and proliferation including *MEF2C, JUND*, and *EBF1*.

**Figure 4.**
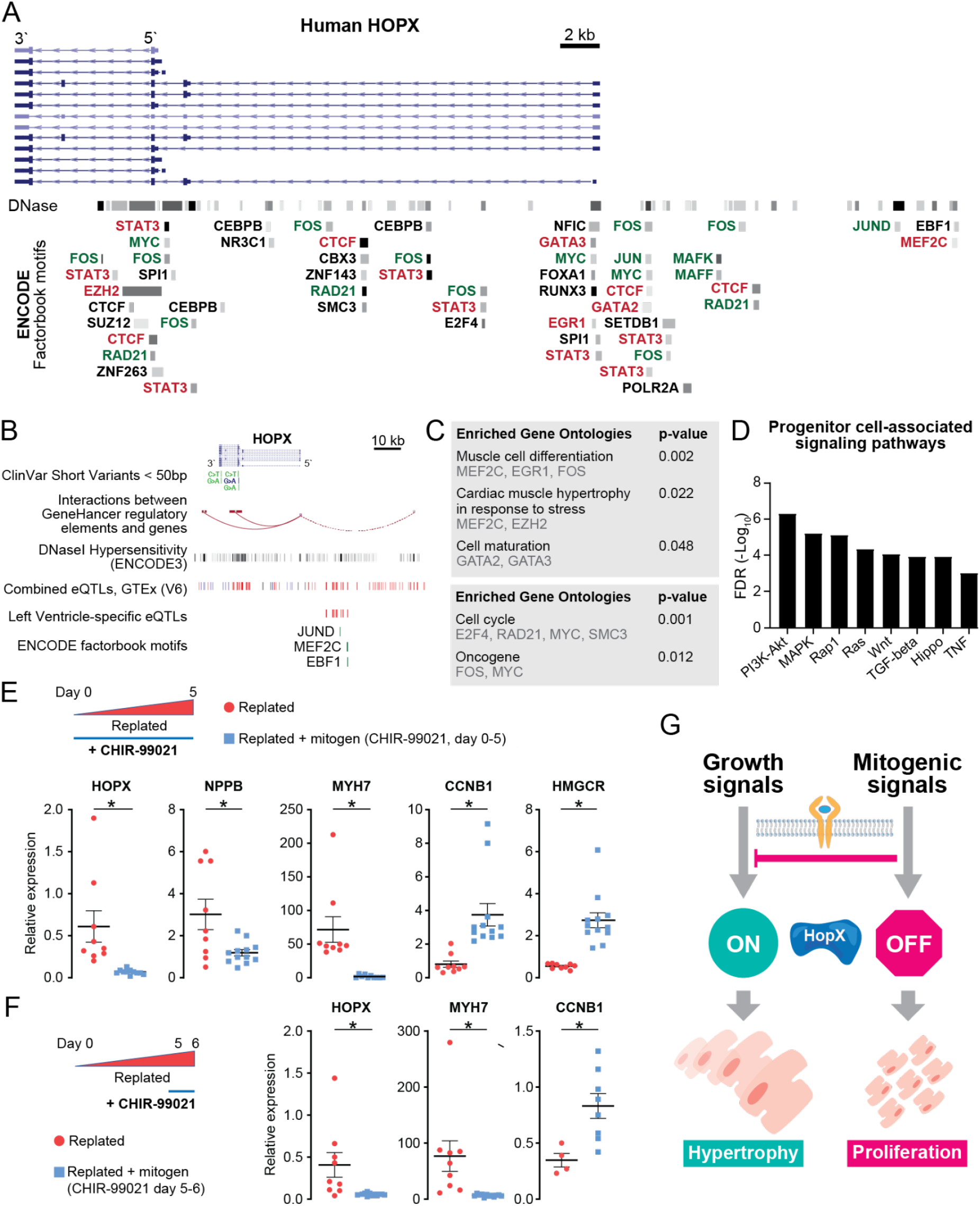
Cell growth and mitogenic signals compete for transcriptional control of *HOPX*. (**A-C**) UCSC genome browser analysis of the human *HOPX* locus identifies diverse *HOPX* transcripts aligned with DNase hypersensitive domains and ENCODE factorbook motifs (**A**), HOPX-associated ClinVar genetic variants, GeneHancer interactions, and left ventricle-specific expression quantitative loci (eQTL) (**B**), and gene ontology analysis of ENCODE Factorbook motifs within 50 kb of the *HOPX* locus (**C**). (**D**) KEGG signaling pathways significantly associated with cardiomyocyte progenitor differentiation *in vivo* (see **Figure 1E-F**) identifies key regulatory determinants of heart growth, differentiation, and proliferation as putative upstream regulators of *HOPX* transcription. (**E-F**) hiPSC-derived cardiomyocytes placed under growth conditions (low-density replating) show robust activation of *HOPX* with concomitant activation of genes involved in cell growth (*NPPB*) and myofibrillar maturation (*MYH7*) in addition to reduction in genes associated with cell cycle (*CCNB1*) and mevalonate pathway activity (*HMGCR*). Cardiomyocytes placed in growth conditions plus mitogenic stimulation (replated + CHIR-99021) continuously during cell growth (days 0-5) (**E**) or for 24 hours (days 5-6) after growth conditions are established (**F**) results in downregulation of *HOPX* and reversal of downstream maturation gene programs. Statistical significance tested using student’s t-test with Welch’s correction. *n > 6* biological replicates per analysis. Error bars show mean ± S.E.M. P<0.01 (*). (**G**) Model for competitive transcriptional regulation of *HOPX* by growth and mitogenic signaling pathways.

To identify putative upstream pathways controlling transcription of *HOPX*, we first analyzed ENCODE Factorbook motifs enriched within 50 Kb of the *HOPX* locus (**Figure 4A-C**). These data show significant enrichment in motifs of cell cycle regulators including members of the p38/MAPK signaling pathway (MYC, JUN, FOS) as well as factors governing cardiac cell growth pathways (EGR1, MEF2C). To identify signaling pathways that may act to regulate the HOPX locus at least in part via these enriched motifs at the HOPX locus, we used KEGG pathway enrichment to analyze cardiac progenitor- and maturation-associated gene programs from *in vivo* time course data (**Figure 1E-H, Figure S1, Table S1**). Cardiac maturation gene programs were enriched for adrenergic signaling, HIF1 signaling, as well as glucagon and oxytocin signaling that control physiological and metabolic characteristics of cardiomyocytes (**Table S1**). In contrast, cardiac progenitor genes were enriched in factors controlling cardiac cell growth and proliferation including Wnt, MAPK, and Hippo pathways (**Figure 4D**).

We functionally tested these cell growth and proliferation pathways as upstream determinants of *HOPX* transcription. hiPSC-derived cardiomyocytes were replated under hypertrophic growth conditions(Friedman et al., 2018) with or without mitogenic stimulation induced by the GSK3β inhibitor small molecule CHIR-99021 (5µM). These data reveal that growth conditions induce *HOPX* transcription concomitant with myofibrillar maturation (e.g. increased expression of *MYH7*) and cell growth (*NPPB*) and downregulation of genes associated with cell proliferation (*CCNB1*) and mevalonate pathway (*HMGCR*) activity. In contrast, growth conditions plus mitogenic stimulation resulted in repression of *HOPX* transcription together with genes associated with cardiac progenitor programs including increased cell cycle and mevalonate pathway activity and reduced cell growth and myofibrillar maturation (**Figure 4E**). We next analyzed whether mitogenic signals could suppress *HOPX* transcription under conditions of established growth stimulation. Cardiomyocytes were placed under conditions to activate cardiac growth and maturation gene programs for 5 days and then exposed to mitogenic stimulation for 24 hours. Indeed, acute addition of a mitogenic stimulus significantly suppressed *HOPX* transcription coupled with induction of cardiac progenitor gene programs (**Figure 4F**), suggesting a dominant regulatory role of mitogenic stimuli in repressing *HOPX* transcription during differentiation.

Collectively, these data provide a model for competitive control of *HOPX* transcription (**Figure 4G**). Cell growth signaling promotes *HOPX* expression to activate maturation gene programs. Mitogenic stimulation such as Wnt signaling activation suppresses *HOPX* expression enabling activation of cardiac progenitor cell gene programs. In competitive assays, mitogenic signals override growth signals to repress *HOPX* transcription.

### *HOPX* controls cardiomyocyte physiological maturation

To determine the functional significance of *HOPX* control of cardiomyocyte gene networks, we used cell-, tissue-, and organ-level assays to evaluate *HOPX* regulation of cardiac physiology. We initially assessed cellular-level phenotypes using high purity iPSC-derived cardiomyocytes (**Figure 5A**) focusing on *HOPX*-regulated biological processes (**Figure 3D-G**). Based on *HOPX* down-regulation of HIF-1 signaling we assessed cardiomyocyte sensitivity to hypoxia using a protocol to model ischemia-reperfusion injury *in vitro*(Redd et al., 2021). hiPSC-CMs were incubated under hypoxic conditions (0.5% O_2_) for 18 hours followed by four hours under normoxic conditions. Consistent with *HOPX-*dependent regulation of HIF-1 signaling, we observed significantly reduced cardiomyocyte sensitivity to acute hypoxic stress in *HOPX* KD compared to controls as measured by release of lactate dehydrogenase (LDH). (**Figure 5B**).

**Figure 5.**
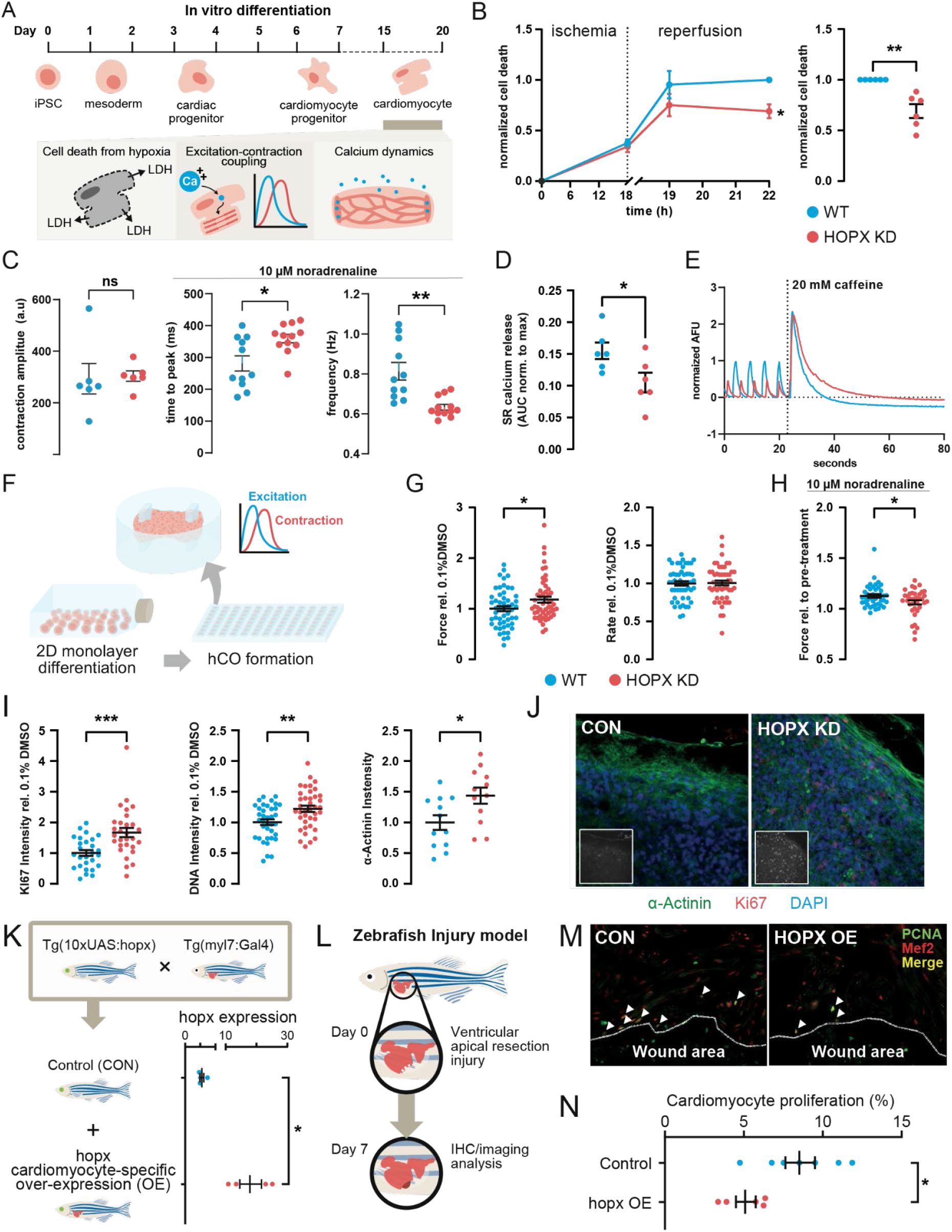
*HOPX* directly regulates cardiomyocyte cell cycling and function with implications in tissue function *in vitro* and *in vivo*. **(A)** Schematic of iPSC cardiac differentiation for assaying HOPX-dependent cardiomyocyte function. (**B-E**) Physiological assays of replated cardiomyocytes using control (CON, no dox treatment) versus HOPX knockdown (KD, dox treatment) cells. **(B)** Cell death (lactate dehydrogenase release) analysis during *in vitro* acute ischemic stress. Left plot shows time course analysis and right plot shows mean data at completion of the assay. **(C)** MUSCLEMOTION contractility analysis of cardiomyocytes under basal conditions and following 10 µM noradrenaline treatment. (**D-E**) Calcium transients measured by the FLIPR Tetra system. **(D)** SR calcium release under basal conditions (average area under the curve for a single transient) normalized to maximum release following 20 mM caffeine stimulation. **(E)** Example calcium trace (AFU = arbitrary fluorescence units) with acute caffeine addition (dotted black line). **(F)** Schematic of experimental strategy to assess proliferation and physiology of hiPSC-derived cardiomyocytes from three-dimensional (3D) cardiac organoids culture. (**G-H**) Analysis of human cardiac organoids reveal *HOPX* KD (DOX) results in significantly increased contractility compared to control (CON) tissues with no change in beat rate (**G**). However, acute addition of 1 µM noradrenaline results in a significant reduction in contractile force with *HOPX* KD vs control (**H**). (**I-J**) Immunostaining analysis revealed increased cardiomyocyte cell cycling measured by Ki67, DNA, and α-actinin intensity in *HOPX* KD compared to control (CON) hCOs as shown in mean data (**I**) and representative images (**J**). **(K)** Schematic of transgenes and experimental strategy to generate zebrafish overexpressing *hopx* in *myl7*^*+*^ myocardium. GFP (green) expression in the eyes driven by an α-crystallin promoter indicates incorporation of *10xUAS:hopx* transgenes, and RFP (red) expression in the heart indicates incorporation of *myl:Gal4* transgenes. Quantitative RT-PCR (qPCR) shows expression of *hopx* relative to housekeeping gene *rpl13* in adult control *Tg(10xUAS:hopx)* and *hopx* overexpressing (OE) *Tg(10xUAS:hopx; myl7:Gal4)* zebrafish hearts (*n* = 4 replicates per condition). **(L)** Schematic of experimental strategy assessing zebrafish cardiomyocyte cycling in response to cardiac ventricular apical resection at seven days post-injury (DPI). (**M-N**) Quantification of cardiomyocyte cycling measured from immunostained sections (**M**) of adult control (CON) and *hopx* OE zebrafish hearts (OE) at seven DPI shown as the percentage of PCNA^+^/Mef2^+^ cells divided by total Mef2^+^ cardiomyocytes in the border zone of the injury (**N**). Representative images are provided from cardiac ventricular sections containing wound areas (outlined in white dotted line) from adult control (left) and *hopx* OE (right) zebrafish hearts at seven DPI immunostained for PCNA (green) and Mef2 (red). Arrowheads indicate cycling cardiomyocytes (PCNA^+^/Mef2^+^). Sample number and statistics: for cell assays (**B-E**) *n* = 4-6 per condition. For organoid assays (**F-J**) *n* = 7-20 tissues per condition. For zebrafish studies (**K-N**) *n* = 7 control and 5 *hopx* OE zebrafish. For all comparisons, statistical significance was tested using student’s t-test with Welch’s correction or two-way mixed ANOVA using Šídák post Hoc test (panel B, left). Error bars show mean ± S.E.M. P<0.05 (*). P<0.01 (**).

*HOPX* KD results in down-regulation of gene programs governing cardiomyocyte excitation-contraction (E-C) coupling and gene programs governing adrenergic responsiveness (**Figure 3D-G**). We therefore assessed *HOPX-*dependent parameters of contractility and calcium handling in iPSC-derived cardiomyocytes at baseline and after stimulation with noradrenaline. *HOPX* KD showed no significant impact on any parameter of cardiomyocyte contraction or relaxation kinetics at baseline. However, in response to 10 µM noradrenaline, *HOPX* KD resulted in significantly slower contraction kinetics and beat rate indicative of an attenuated adrenergic response compared to robust inotropic stimulation observed in WT cells (**Figure 5C** and **Figure S7**). Total sarcoplasmic reticulum (SR) calcium load was similar between control and *HOPX* KD cardiomyocytes as measured by peak calcium release after exposure to 20 mM caffeine (**Figure 5D-E**). However, HOPX knockdown resulted in a significant reduction in the ratio of baseline calcium transient peak normalized to peak caffeine-induced calcium release (**Figure 5D-E**). This suggests that while *HOPX* KD does not impact total calcium storage, *HOPX* KD cardiomyocytes have reduced beat-to-beat SR calcium release.

### *HOPX* increases force generating units in iPSC-derived organoids

Tissue-level force was assessed in immature human cardiac organoids (hCO) consistent with the developmental activity of *HOPX* (Mills et al., 2019) (**Figure 5F-H**). *HOPX* KD significantly increased force with no impact on beat rate compared to controls (**Figure 5G**). Loss of *HOPX* also significantly reduced the increases in response to adrenergic stimulation by 1 µM noradrenaline (**Figure 5H**). As these findings are consistent with a greater number of immature cardiomyocytes, we hypothesized that cardiac progenitor programs induced by *HOPX* loss-of-function results in proliferation. Indeed, immunohistochemical analysis of hCOs revealed significantly increased cardiomyocyte proliferative activity in *HOPX* KD tissues with increased staining intensity of Ki-67, DNA, and α-actinin (**Figure 5I-J**).

### Zebrafish hopx inhibits cardiomyocyte proliferation in whole organs *in vivo*

While human hearts lack endogenous cardiomyocyte proliferative capability, zebrafish hearts naturally regenerate after injury or resection through proliferation of endogenous cardiomyocytes(Jopling et al., 2010; Kikuchi et al., 2010). We therefore tested whether *hopx* over-expression could repress innate cardiomyocyte proliferation *in vivo* during the acute regenerative phase in a zebrafish model of apical resection(Poss et al., 2002; Vivien et al., 2016). We used the Gal4:UAS transactivation system to overexpress *hopx* in a cardiomyocyte-specific manner. Stable transgenic zebrafish lines (*Tg(10xUAS:hopx)* were crossed with a cardiomyocyte-specific Gal4 driver line (*Tg(myl7:Gal4)*) to generate progeny that show robust overexpression of *hopx* in cardiomyocytes (OE; *Tg(10xUAS:hopx; myl7:Gal4)* compared to siblings (CON) (**Figure 5K**). Analysis of gene expression and contractility during homeostasis at 5 days post fertilization (DPF) revealed no significant differences in cardiac physiology in *hopx* OE zebrafish versus siblings (**Figure S8I**).

Control and *hopx* OE adult zebrafish were subjected to a resection model of cardiac injury whereby approximately 20% of the apical ventricle was surgically removed (**Figure 5L**). Hearts were extracted at 7 days post injury (DPI). The border zone adjacent to the wound area was analyzed by immunohistochemistry for expression of PCNA^+^ and Mef2^+^ proliferating cardiomyocytes. This revealed significantly reduced cardiomyocyte proliferation in *hopx* OE zebrafish compared to controls (40.15% reduction) (**Figure 5M-N**). Collectively, these data provide comprehensive profiling of cell, tissue, and organ-level physiology to link *HOPX-* regulated molecular programs with cardiomyocyte physiology.

### Gene networks governed by HOPX are associated with heart structure and function traits

We analyzed gene knockout phenotype databases(McLean et al., 2010) and found enrichment of HOPX-bound loci (from DamID-seq: 3,467 genes) associated with cardiac structure and function phenotypes including cardiac contractility, cardiac fiber size, responses to myocardial infarction, exercise endurance, and sarcomere morphology (**Figure S9A**). We therefore tested the hypothesis that human *HOPX* genetic variants influence natural variation in cardiac complex trait phenotypes. Analysis was performed using genome-wide association studies (GWAS) linking cardiac MRI structure and function data to patient DNA variants from UKBiobank(Pirruccello et al., 2020). While left ventricular-specific eQTLs overlap motifs of key regulators of cardiac maturation and proliferation (**Figure 4B**), variant proximity analysis or eQTL analysis (Summary Mendelian Randomisation) indicate that genetic variation at the *HOPX* locus alone does not significantly influence cardiac structural and functional traits (**Figure S9B**). Indeed, constraint metrics measured by gnomAD using the ensemble canonical transcript (ENST00000554144.1) indicate that the *HOPX* locus is not significantly intolerant to genetic variation (predicted loss of function, pLoF o/e = 1.35).

We next tested whether HOPX bound (3,467 genes by DamID-seq) and/or genes regulated by HOPX (650 CRISPRi DE genes by CAGE-seq) influence natural variation in human cardiac organ development *in vivo*. First, we used fastBAT to analyze the aggregated effects of genetic variants ± 50 kb of a gene, accounting for linkage disequilibrium between genetic variants. Indeed, genes regulated by *HOPX* including *MYH6/7* and *ALPK3* among others, were significantly associated with variation in parameters including left ventricular systolic and diastolic volumes and ejection fraction (**Table 1**). Moreover, *HOPX* DE genes were significantly over-represented in genes regulating left ventricular end systolic volume (LVESV_BSI, P<0.05 Fisher’s Exact Test).

**Table 1.** eQTL-based analysis of genetic associations with cardiac structure and function using UKBiobank GWAS data performed using Summary Mendelian Randomization. Chi-squared or Fisher’s test was used to calculate over-representation of HOPX bound genes among genes significantly associated with a cardiac trait. * P < 0.05 by Fisher’s test was used to calculate over-representation of DE genes among genes significantly associated with a cardiac trait. Left ventricular end systolic volume (LVESV), left ventricular end diastolic volume (LVEDV), left ventricular ejection fraction (LVEF), body surface area indexed (BSI). HOPX DE indicates which direction the gene is differentially expressed with HOPX KD.

| *Gene* | **LVESV** | **LVESV_**<br>**BSI (\*DE)** | **LVEDV** | **LVEDV_**<br>**BSI** | **LVEF** | **HOPX DE** |
| --- | --- | --- | --- | --- | --- | --- |
| **AGO2** | 2.82E-07 | 4.08E-08 |  |  | 5.93E-07 | Down |
| **AKR1A**<br>**1** | 3.95E-08 | 6.24E-08 | 1.49E-07 |  |  | Down |
| **ALPK3** | 5.46E-11 | 4.63E-10 |  |  | 6.13E-10 | Down |
| **GPR15**<br>**3** | 2.18E-07 | 5.46E-07 |  |  |  | Down |
| **MYH7** |  |  | 7.42E-07 |  |  | Down |
| **RGS10** | 8.7E-10 | 1.87E-10 | 2.37E-07 | 3.51E-09 |  | Down |
| **SPON1** |  | 3.82E-07 |  |  |  | Down |
| **TESK2** | 5.11E-07 |  | 7.22E-07 |  |  | Down |
| **MYH6** |  |  | 4.28E-07 |  |  | Up |
| **PKN2** |  |  |  |  | 2.94E-07 | Up |

Second, we used summary-data-based Mendelian Randomization (SMR) to evaluate the association between changes in gene expression of genes bound or regulated by *HOPX* and cardiac phenotypes (**Table 2**). We found that genes with eQTLs significantly associated with left ventricular chamber structural and functional traits were significantly over-enriched for *HOPX* bound genes for LV end systolic volume (LVESV and LVESV_BSI, P<0.05 Chi Square Test) and *HOPX* DE genes for LV end diastolic volume (LVEDV_BSI, P<0.05 Fisher’s Exact Test). Additionally, we identified direct targets of *HOPX* (*SPON1, SPATA24, PRKCA, MTSS1*) to be significantly associated with left ventricular chamber dimensions and function. These data indicate that *HOPX* governs an unexpectedly high number of genes influencing human cardiac structure and function complex trait phenotypes.

**Table 2.** Summary Mendelian Randomization providing eQTL analysis of genetic associations with cardiac structure and function using UKBiobank GWAS data. * P < 0.05 by Fisher’s Exact Test. # P < 0.05 Chi Square Test. Left ventricular end systolic volume (LVESV), left ventricular end diastolic volume (LVEDV), left ventricular ejection fraction (LVEF), body surface area indexed (BSI). HOPX DE indicates which direction the gene is differentially expressed with HOPX KD. HOPX locus indicates whether HOPX is identified at that locus from DamID data.

| *Gene* | **LVESV**<br>**(#, bound)** | **LVESV_BSI**<br>**(#, bound)** | **LVEDV** | **LVEDV_**<br>**BSI (\*DE)** | **LVEF** | **HOPX**<br>**locus** | **HOP**<br>**X KD** |
| --- | --- | --- | --- | --- | --- | --- | --- |
| **WDR73** | 3.92E-06 | 6.39E-06 |  |  | 7.72E-06 |  |  |
| **SPON1** | 4.03E-06 | 6.71E-07 |  |  |  | bound |  |
| **SPATA24** |  |  |  |  | 2.76E-06 | bound |  |
| **SMARCB1** | 1.4E-06 | 3.11E-07 |  |  | 5.48E-06 |  |  |
| **PRKCA** | 1.59E-06 | 2.33E-07 |  |  | 1.89E-06 | bound |  |
| **MTSS1** | 1.22E-09 | 1.09E-08 | 2.08E-08 | 2.98E-07 | 9.03E-07 | bound | DE |
| **MMP11** | 3.86E-08 | 1.98E-08 |  | 5.68E-06 | 3.78E-08 |  |  |
| **MLF1** |  | 6.86E-06 |  |  |  |  |  |
| **MAPT** | 5.4E-08 | 2.45E-07 |  | 9.6E-07 |  |  |  |
| **LMF1** |  | 5.29E-07 |  |  | 5.37E-06 |  |  |
| **LINC00964** | 1.44E-06 |  | 4.62E-06 | 3.4E-06 |  |  |  |
| **FKBP7** | 1.31E-06 | 2.81E-06 | 5.68E-06 |  |  |  |  |
| **EXOSC5** |  |  | 2.65E-06 |  |  |  |  |
| **DNAJC18** |  | 3.53E-06 |  |  | 1.03E-06 |  |  |
| **CLCNKA** | 3.94E-06 | 3.21E-06 |  |  | 4.77E-07 |  |  |
| **AC103965.1** | 7.48E-06 |  |  |  |  |  |  |

## DISCUSSION

Mechanisms governing cardiomyocyte progenitor and maturation gene programs remain a focus point for understanding genetic causes of heart disease and identifying strategies for cardiac regeneration. While HOPX has garnered significant attention in the biological mechanisms of heart development(Chen et al., 2002; Trivedi et al., 2010), its role in diverse developmental contexts(Palpant et al., 2017a) and cancer(Liu and Zhang, 2020) reinforce its underlying role underpinning cell identity and differentiation in health and disease. This study deconstructs genetic mechanisms of HOPX regulation of cardiomyocyte differentiation, positioning it as a molecular switch governing cardiac progenitor and maturation gene networks. Despite the inability of HOPX to directly bind DNA, we show that HOPX interacts with and regulates core cardiac developmental gene programs controlling cardiomyocyte cell metabolism, signaling, electromechanical coupling, and transcriptional regulation that underpin molecular and functional cell state changes.

Mechanisms controlling cell growth and proliferation are central to the biology of organ morphogenesis and function in health and disease. This study provides a detailed mechanistic basis for *HOPX* as a central determinant of these processes in cardiomyocytes, with evidence linking molecular control of cardiac developmental gene programs to functional phenotypes via regulation of *HOPX* transcription. We show that upstream mitogenic and hypertrophic signaling pathways compete for activation of progenitor and maturation gene programs by transcriptional repression or activation of *HOPX*, respectively. Mitogenic stimuli override cell growth pathways to suppress *HOPX* expression early in differentiation thereby promoting progenitor cell proliferation. While we show that Wnt agonists act as mitogenic stimuli to potently repress transcription of *HOPX*, Wnt activation reduces force production(Mills et al., 2019) thereby suggesting alternative signaling pathways governing *HOPX* transcription in cardiomyocytes. Studies in cancer indicate that HOPX is a tumor suppressor, acting via a HOPX-Ras-MAPK network to determine cell senescence. Indeed, *HOPX* loss-of-function in iPSC-derived hCOs phenocopies MAPK signaling inhibition(Mills et al., 2017a), suggesting a regulatory relationship between MAPK signaling as a potential added mechanism for controlling *HOPX* transcription during cardiomyocyte differentiation.

The potent molecular control mechanisms governing *HOPX* transcription are reinforced by evidence positioning *HOPX* as a mediator of two mutually exclusive heart development gene programs underpinning progenitor and maturation cell states. We show that *HOPX* binds to and is associated with binary on/off regulation of entire signaling pathways, potent transcription factors governing heart field specification, myofilament isoform changes, ion channels controlling cardiomyocytes E-C coupling kinetics, cell stress pathway mediators, and metabolic programs that are well-established features of cardiomyocyte maturation(van Hengel et al., 2017; Mills et al., 2019). We demonstrate that these *HOPX* dependent gene programs translate functionally into physiological cell state changes across diverse cardiac myocyte functional phenotypes including cell stress responses, electro-mechanical coupling, and tissue-level force. Unlike many mitogenic stimuli that reduce force production(Mills et al., 2017a), we show that *HOPX* depletion in cardiac progenitor cells increases tissue-level contractility through proliferation of immature force generating units. To promote functional maturation of cardiomyocytes, hypertrophic cell signaling pathways act via *HOPX* to repress progenitor gene programs and up-regulate structural, electromechanical, and metabolic programs.

Natural genetic variation underpins fundamental changes in complex traits, providing powerful methods to predict disease risk and identify targets more likely to succeed as next-generation therapeutics(King et al., 2019). Despite our studies showing potent effects of HOPX transcriptional abundance, natural human genetic variation at the *HOPX* locus or *HOPX*-associated eQTLs do not significantly influence cardiac structure and function in the human population based on statistical power of available cohort sizes. Indeed, *HOPX* variants are only significantly associated with a small number of blood phenotypes with no links to disease in population genetic data and no reported pathogenic variants reported in ClinVAR. However, drawing on our detailed analysis of the *HOPX* regulome, we show that *HOPX-*dependent gene programs are more likely to determine phenotypic variation in heart structure and function than would be expected by chance. This further reinforces the critical function of *HOPX* in fundamental regulation of cardiac organ development, structure, and function.

Taken together, this study positions *HOPX* in the context of upstream signaling pathways and downstream gene programs governing cardiomyocyte proliferation and maturation at molecular and physiological levels. The results establish a mechanistic basis for *HOPX* as a primary determinant of heart development, providing insights linking *HOPX* to gene programs, signaling pathways, and physiological functions that govern the switch between mutually exclusive progenitor and maturation cell states. These insights not only establish a molecular mechanism for *HOPX* in cardiomyocyte differentiation, but also provide a robust basis for therapeutic strategies targeting *HOPX* and its associated signaling pathways and gene programs.

## ACKNOWLEDGMENTS

We thank the University of Queensland Genome Innovation Hub for support of CAGE-seq analysis. SWC and GJF are supported by funding from the Australian National Health and Medical Research Council (NHMRC) (GNT1173711 to GJF and GNT1161832 to SWC) and by the Mater Foundation (Equity Trustees / AE Hingeley Trust). This work has been supported by grant funding to NJP (ARC SR1101002, DP170101217, NHMRC APP1143163, and National Heart Foundation FLF101889). RPH was supported by grants from NHMRC (2008743, 1118576, 1061539). We would like to thank Christine Seidman for mouse embryonic heart development single cell data(DeLaughter et al., 2016).

## AUTHOR CONTRIBUTIONS

**CEF**. Engineered *HOPX* CRISPRi cells and hopx OE zebrafish, designed HOPX DamID vectors, performed all stem cell differentiation assays for *HOPX* loss-of-function, CAGE-seq, and DamID-seq assays, performed analysis of baseline hopx OE zebrafish and gene expression, coordinated collaborative requirements for completion of the study, and wrote the manuscript. **SWC and GJF**. Prepared the DamID-seq libraries, performed analysis of HOPX DamID data and integration with CAGE-seq data and wrote the manuscript. **RJM, HV, and JEH**. Performed cardiac organoid assays. **MO and KK**. Performed zebrafish apical resection assays. **HSC and XC**. Assisted with completion of stem cell differentiation and DamID-seq assays and performed assays related to Figure 4E-G. **SS and YS**. Performed bioinformatics analysis of single cell RNA-seq (SS and YS) and CAGE data (SS). **DM**. Performed GWAS analysis. **MAR**. Performed *HOPX* loss-of-function hypoxia sensitivity assay and cardiomyocyte electromechanical coupling analyses (contractility and calcium). **RB and RPH**. Assisted in design of HOPX DamID and generated DamID lentiviruses. **SP and BMH**. Generated hopx OE zebrafish and assisted with zebrafish colony maintenance, genotyping, and analysis. **JDA and KAS**. Embedded zebrafish in agarose and recorded videos for cardiac physiology assessment. **SA, SY, and QN**. Performed CAGE-sequencing and assisted with analysis. **NJP**. Supervised the project, designed experiments, analyzed data, raised funding to complete the work, and wrote the manuscript.

## METHODS

### Experimental model and subject details

The authors will make the data, methods used in the analysis, and materials used to conduct the research available to any researcher for purposes of reproducing the results or replicating the procedure. *Study Design*. All animal and cell experiments were performed in accordance with protocols approved by the Animal and Human Ethics Committee of The University of Queensland (UQ) (IMB/171/18 and IMB/030/16/NHMRC/ARCHF). All animals used in this study received humane care in compliance with Australian National Health and Medical Research Council guidelines and the *Guide for the Care and Use of Laboratory Animals* (U.S. National Institutes of Health). All hiPSC studies were performed with consent from the UQ Institutional Human Research Ethics approval (HREC #2015001434). Additional information for each set of experiments, such as statistical analyses, exclusion criteria, and sample size, are detailed in the relevant methods section or figure legend.

### Analyses of Human and Mouse Embryonic Development at Single-Cell Resolution

We integrated mouse time course heart development single-cell datasets(DeLaughter et al., 2016; Lescroart et al., 2018; Li et al., 2016) and human time course single-cell dataset comprised of RNA-seq(Asp et al., 2019) and two single nucleus datasets(Nicin et al., 2021; Sim et al., 2021). Annotations and time points were defined based on the original publication. Data integration was performed using standard parameters in *Seurat* (version 3.0.0)(Stuart et al., 2019). We performed PCA and computed 50 principal components then performed UMAP dimensionality reduction. **Data availability**: Data were obtained from Gene Expression Omnibus (GEO) under accession number GSE76118(Li et al., 2016) and GSE100471(Lescroart et al., 2018). Data for (DeLaughter et al., 2016) were kindly provided by the authors. Data for (Asp et al., 2019) were obtained from https://www.spatialresearch.org. Data from (Sim et al., 2021) and (Nicin et al., 2021) were kindly provided by the authors.

### Single Cell Gene Expression Correlation Analyses

The Pearson correlation was calculated for select progenitor and maturation genes compared to all expressed genes in every individual cell comprising the integrated developmental mouse heart scRNA-seq dataset(DeLaughter et al., 2016; Lescroart et al., 2018; Li et al., 2016). Scatter plot panels were generated using the ggplot2 (v3.3.2) R (v3.6.0) package. The scatter plots illustrate the expression of select genes in each cell with the cells ordered by the reference gene expression level.

### Generation and Maintenance of Human ESC/iPSC Lines

All human pluripotent stem cell studies were carried out in accordance with consent from the University of Queensland’s Institutional Human Research Ethics approval (HREC#: 2015001434). WTC CRISPRi GCaMP hiPSCs (Karyotype: 46, XY; RRID CVCL_VM38), generously provided by M. Mandegar and B. Conklin (UCSF, Gladstone Institute), was generated using a previously described protocol(Mandegar et al., 2016). WTC CRISPRi *HOPX* g1 (XY; RRID CVCL_VQ27) hiPSCs were generated in a previous study (Friedman et al., 2018). All cells were maintained as previously described with slight adaptions(Friedman et al., 2018; Palpant et al., 2017b). Briefly, cells were maintained in mTeSR Plus media with supplement (Stem Cell Technologies, Cat.#05825) at 37°C with 5% CO_2_. All WTC CRISPRi and WTC CRISPRi *HOPX* g1 hiPSC lines were maintained on Vitronectin XF (Stem Cell Technologies, Cat.#07180) coated plates.

### hiPSC Cardiac Differentiation

Cardiac-directed differentiation using a monolayer platform was performed with a modified protocol based on previous reports(Burridge et al., 2014; Friedman et al., 2018; Lian et al., 2012; Palpant et al., 2017b). On day ™1 of differentiation, hiPSCs were dissociated using 0.5% EDTA, plated onto Vitronectin XF coated plates at a density of 1.8 × 10^5^ cells/cm^2^, and cultured overnight in mTeSR Plus media. Differentiation was induced on day 0 by first washing with PBS, then changing the culture media to RPMI (ThermoFisher, Cat.#11875119) containing 3 μM CHIR99021 (Stem Cell Technologies, Cat.#72054), 500 μg/mL bovine serum albumin (BSA, Sigma Aldrich, Cat.#A9418), and 213 μg/mL ascorbic acid (Sigma Aldrich, Cat.#A8960). After 3 days of culture, the media was replaced with RPMI containing 500 μg/mL BSA, 213 μg/mL ascorbic acid, and 1 μM XAV-939 (Stem Cell Technologies, Cat.#72674). On day 5, the media was exchanged for RPMI containing 500 μg/mL BSA, and 213 μg/mL ascorbic acid without any additional supplements. From day 7 onward, the cultures were fed every 2 days with RPMI containing 2% B27 supplement plus insulin (Life Technologies Australia, Cat.#17504001). Cells were dissociated on day 15 of differentiation using 0.5% EDTA, stopped with 1:1 FBS in DMEM:F12, filtered with a 100 μm strainer, replated at a density of 1.58 × 10^5^ cells/cm^2^ in Vitronectin XF-coated 24 well plate into cardiomyocyte media (RPMI, 2% B27 supplement plus insulin [Life Technologies Australia, Cat.#17504001], 10% FBS, and 10 µM Y-27362 ROCK inhibitor [Stem Cell Technologies, Cat.#72308]) and cultured overnight. FBS and ROCK inhibitor-containing media was replaced the following day with standard media and replaced every 48 hours thereafter.

### DamID-Sequencing

**Lentiviral Transduction of hiPSC-Derived Cardiomyocytes:** One day in advance of lentiviral transduction, hiPSC-derived cardiomyocytes (15 days post-activation) were replated into a 6wp at 2.083 × 10^4^ cells/cm^2^. Day 16 cardiomyocytes were transduced one day after replating using spinfection with polybrene and incubated in transduction media for 72 hours. Day 19 cardiomyocytes were washed thrice with PBS and harvested for genomic DNA isolation. **DamID-seq Library Preparation and Sequencing:** DamID-sequencing was performed as previously described(Marshall et al., 2016). Briefly, genomic DNA from four independent replicates of HOPX-Dam and corresponding Dam-alone was extracted, digested with *Dpn*I (NEB) to cleave methylated GATC sites and adaptors were ligated. The DNA was then digested with *Dpn*II to remove fragments containing unmethylated GATC sites. The DNA was PCR amplified using the MyTaq polymerase (Bioline). DNA was sonicated to an average size of 300bp and adaptors were removed by *Alw*I (NEB) digestion. 1µg of sonicated DNA was prepared for sequencing using the NEBNext Ultra™ II DNA Library Prep Kit for Illumina without size selection and amplified with 3 cycles of PCR. **DamID-seq Analysis:** Four independent replicates of HOPX-Dam with corresponding Dam-alone controls were sequenced on the Illumina NovaSeq instrument, generating a median of 30.6 million 100bp single-end reads per sample. Reads were aligned to the human genome (hg38) using bowtie2(Langmead and Salzberg 2012). Reads were binned into GATC fragments and normalized to the corresponding Dam control using the damidseq_pipeline (Marshall and Brand, 2015). Bedgraph files were converted to bigwig format using the bamCoverage function of deepTools(Ramírez et al., 2014). Correlations between replicates were calculated across 10kb genomic bins using the deepTools multiBigwigSummary and plotCorrelation tools (Ramírez et al., 2014). An outlier (replicate 4) was identified and excluded from further analyses. The remaining three replicates were quantile normalized and peaks were identified with a DamID peak caller with the specification –frac=500000, FDR<0.01 (https://github.com/owenjm/find_peaks,(Marshall and Brand, 2015)). Consensus peaks between the three individual replicates and the average of the replicates were determined using BEDTools intersectBed (Quinlan and Hall, 2010). Gene ontology enrichment of peaks was performed using Genomic Region Enrichment of Annotation Tools (GREAT, (McLean et al., 2010)). **Transcription Factor Binding Site Motif Analysis of HOPX-Dam Consensus Peaks:** Sequences underlying consensus HOPX-Dam peaks were extracted using BEDTools getfasta(Quinlan and Hall, 2010). Sequences were uploaded to the web portal of MEME-ChIP(Machanick and Bailey, 2011) and enriched motifs detected using default settings.

### CAGE-Sequencing

**Library Preparation:** Library preparation was performed by the Genome Innovation Hub (GIH) and UQ Sequencing Facility at the Institute for Molecular Bioscience using the DNAFORM commercial CAGE preparation user guide with the addition of a qPCR quantification step to normalized input for pooling before second strand synthesis **Sequencing:** Sequencing with 5% PhiX spike-in was performed on an Illumina NextSeq500 using the following configurations: Read1 – 76bp, Index1 – 6bp. Loading concentrations were 1.6pM for the HOPX g1 CRISPRi samples, using a modified denaturation and dilution protocol for low-concentration libraries. **TSS Analysis:** CAGE-seq data was analyzed using the CAGEr pipeline(Haberle et al., 2015). **Differential Expression Analysis:** Promoters and genes were assessed for differential expression using DESeq2 described(Love et al., 2014). **Gene Ontology Analysis:** Gene ontology (GO) analysis was performed using DAVID with significance threshold set at FDR < 0.05 with only level 5 biological process, molecular function, and cellular components GO and KEGG terms selected (Huang et al., 2009).

### hiPSC-CM physiological assays in 2D

Cardiomyocytes derived from WTC CRISPRi HOPX g1 cell line (wildtype control: no Dox treatment; HOPX KD: Dox treatment from D0+) were replated on day 15 – 17 of differentiation in 96 well plate format (VTN-coated) at a density of 1.2 × 10^5^ cells/cm^2^ (contractility) or 6.2 × 10^4^ cells/ cm^2^ (ischemia-reperfusion injury), or in 384 well plate format (Corning CellBIND plate, black with clear bottom) at a density of 3.2 × 10^5^ cells/cm^2^. Cells were maintained for an additional 7 days in RPMI supplemented with B27, with the medium replaced every other day, followed by one of the following endpoint assays. **Ischemia-Reperfusion Injury:** To simulate ischemic injury *in vitro*, replated cells were incubated for 18h under hypoxic (0.5% O_2_; 5% CO_2_) culture conditions with 1x Hanks Buffered Sodium Salt (HBSS) (Sigma, without sodium bicarbonate) medium with 12 mM HEPES (Sigma) and pH adjusted to 7.4(Sala et al., 2018). The medium was replaced with RPMI + B27 and cultured for an additional 4 hours under normoxic conditions (∼18.5% O_2_; 5% CO_2_). Supernatant was collected after 18h hypoxia, and after 1- and 4-hours of reoxygenation, and lactate dehydrogenase (LDH) levels were measured using a cytotoxicity detection kit (Roche). Normalized absorbance (Abs) values (Abs at 492 nm minus Abs at 690 nm) were measured using a microplate reader (Tecan). Abs for each well was converted to percent cell death using average absorbance values from low (blank, media only) and high (cells treated with 1% Triton X-100 in RPMI + B27) controls: percent cell death = 100 x [Abs (sample) – Abs (low control)] / [Abs (high control) – Abs (low control)]. Reoxygenation time points represent the cumulative cell death (i.e. cell death after hypoxia plus cell death after reoxygenation). **Contractility:** Brightfield videos (AVI) of cardiomyocyte contractions were acquired at 60 frames per second and analyzed with MUSCLEMOTION ImageJ plugin (Sala et al., 2018). After baseline recording, noradrenaline (final concentration: 10 µM, 1 µM, or 0.1 µM[MR5]) or RPMI + B27 (vehicle control) was added to each well. Cells were incubated for 20 minutes at 37°C, followed by a second video acquisition. **Calcium Imaging:** Cardiomyocytes were incubated with 1x FLIPR Calcium 4 dye (Molecular Devices) diluted in RPMI + B27 for 1.5 h at 37°C. Calcium transients were measured on FLIPR Tetra fluorescent plate reader (Molecular Devices) with excitation wavelengths at 470–495 nm and emission at 515–575 nm. Drug additions were added to each well at t = 20s (acute addition with real time calcium imaging) to yield a final concentration of 20 mM caffeine or dose response of noradrenaline (highest concentration: 100 µM, lowest concentration: 0.1 pM). Cells were incubated for 10 minutes at 37°C, followed by a second acquisition. All data were acquired at 0.1 seconds per read and expressed as normalized arbitrary fluorescence units. Calcium amplitude, maximum calcium, and area under the curve were analyzed post addition (acute and after 10-minute incubation) using ScreenWorks software (Molecular Devices) and normalized to baseline measurements, or as described in the relevant figure legend. Spontaneous beat rate was determined using custom Matlab script (available upon request).

### Quantitative Real-Time PCR

For quantitative real-time PCR was performed as previously described(Palpant et al., 2013). Briefly, total RNA was isolated using the RNeasy Mini kit (QIAGEN, Cat.#74106). First-strand cDNA synthesis was generated using the Superscript III First Strand Synthesis System (ThermoFisher, Cat.#18080051). Quantitative real-time PCR was performed using SYBR Green PCR Master Mix (ThermoFisher, Cat.#4312704) on a ViiA 7 Real-Time PCR System (Applied Biosystems). **qRT-PCR Primers (targeting human genes unless otherwise noted)**:

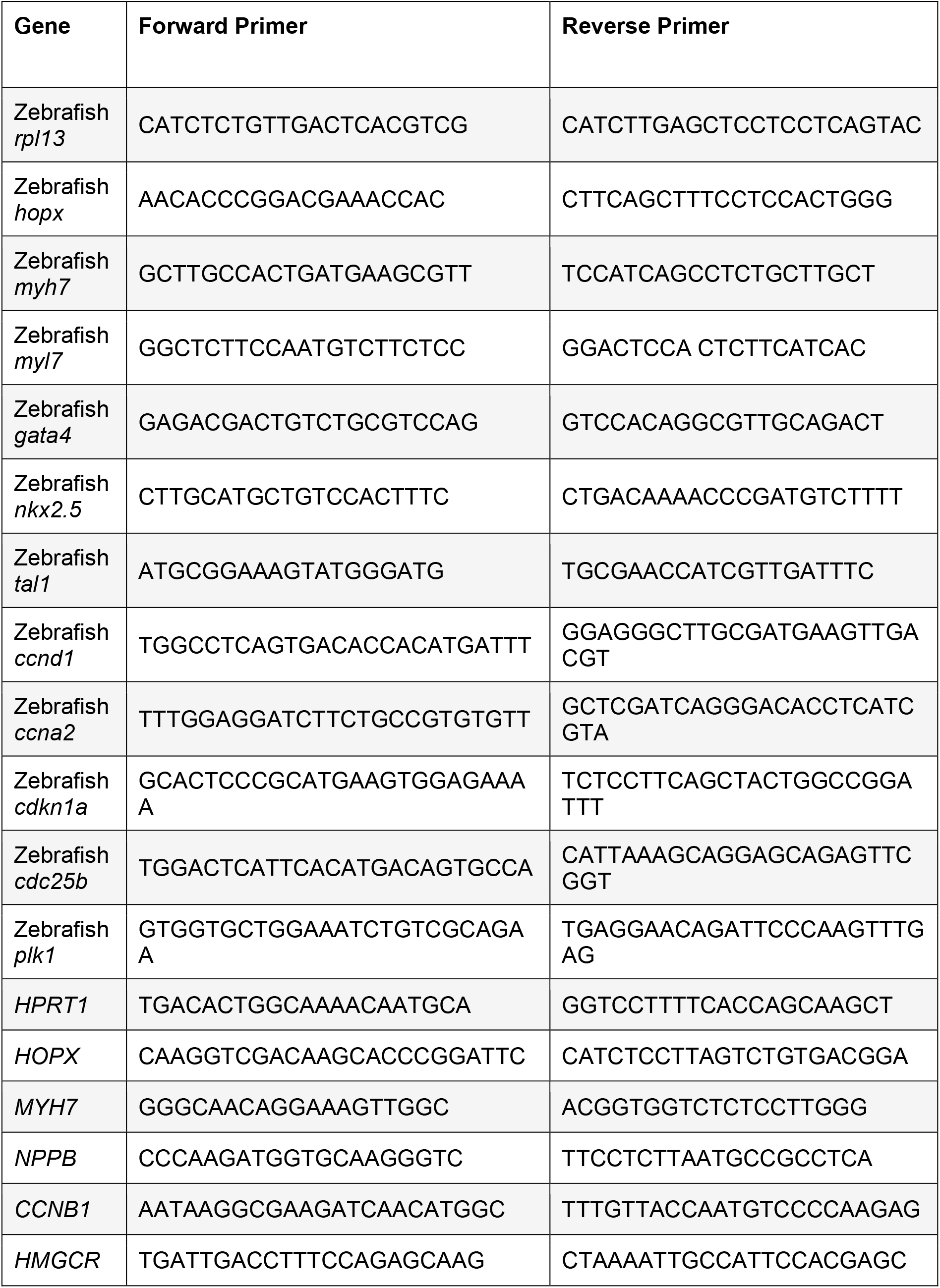

### Heart-Dyno Human Cardiac Organoids (hCO)

**Cardiac Organoid iPSC Differentiation:** Cardiac cells were produced using protocols where cardiomyocytes and stromal cells are produced in the same differentiation culture(Hudson et al., 2012; Mills et al., 2017a, 2017b; Voges et al., 2017); multi-cellular cultures are critical for function(Hudson and Zimmermann, 2011; Tiburcy et al., 2017). hiPSCs were seeded at 2 × 10^4^ cells/cm^2^ in Matrigel-coated flasks and cultured for 4 days using mTeSR-1. They were then differentiated into cardiac mesoderm using RPMI B27-medium (RPMI1640 GlutaMAX+ 2% B27 supplement without insulin, 200 µM L-ascorbic acid 2-phosphate sesquimagnesium salt hydrate [Sigma] and 1% Penicillin/Streptomycin [all ThermoFisher Scientific unless otherwise indicated]) containing 5 ng/mL BMP-4 (RnD Systems), 9 ng/mL Activin A (RnD Systems), 5 ng/mL FGF-2 (RnD Systems) and 1 µM CHIR99021 (Stem Cell Technologies) with daily medium exchange for 3 days. Subsequently, they were specified into a hiPSC-CM/stromal cell mixture using RPMI B27-containing 5 µM IWP-4 (Stem Cell Technologies) followed by another 7 days of RPMI B27+ (RPMI1640 GlutaMAX + 2% B27 supplement with insulin, 200 µM L-ascorbic acid 2-phosphate sesquimagnesium salt hydrate and 1% Penicillin/Streptomycin) with medium exchange every 2-3 days. The differentiated cells were then cultured in RPMI B27+ until digestion at 15 days using 0.2% collagenase type I (Sigma) in 20% fetal bovine serum (FBS) in PBS (with Ca^2+^ and Mg^2+^) for 60 minutes at 37ºC, followed by 0.25% trypsin-EDTA for 10 minutes. The cells were filtered using a 100 µm mesh cell strainer (BD Biosciences), centrifuged at 300 × g for 3 minutes, and resuspended at the required density in CNTL medium: α-MEM GlutaMAX, 10% FBS, 200 µM L-ascorbic acid 2-phosphate sesquimagnesium salt hydrate and 1% Penicillin/Streptomycin. **hCO Fabrication:** Heart-dyno culture inserts were fabricated using standard SU-8 photolithography and PDMS molding practices (Mills et al., 2017a). CNTL medium: α-MEM GlutaMAX (ThermoFisher Scientific), 10% fetal bovine serum (FBS) (ThermoFisher Scientific), 200 µM L-ascorbic acid 2-phosphate sesquimagnesium salt hydrate (Sigma) and 1% Penicillin/Streptomycin (ThermoFisher Scientific). For each hCO, 5 × 10^4^ cardiac cells in CNRL medium were mixed with collagen I to make a 3.5 µL final solution containing 2.6 mg/mL collagen I and 9% Matrigel. The bovine acid-solubilized collagen I (Devro) was first salt balanced and pH neutralized using 10X DMEM and 0.1 M NaOH, respectively, prior to mixing with Matrigel and cells. The mixture was prepared on ice and pipetted into the Heart-Dyno. The Heart-Dyno was then centrifuged at 100 × g for 10 seconds to ensure the hCO form halfway up the posts. The mixture was then gelled at 37ºC for 60 minutes prior to the addition of CNTL medium to cover the tissues (150 µL/hCO). The Heart-Dyno design facilitates the self-formation of tissues around in-built PDMS exercise poles (designed to deform ∼0.07 µm/µN). The medium was changed every 2-3 days (150 µL/hCO). hCOs were cultured in CNTL medium for formation for 5 days and then either kept in CNTL medium culture or changed to maturation medium(Mills et al., 2017a) comprising DMEM (without glucose, glutamine, and phenol red) (ThermoFisher Scientific) supplemented with 4% B27 (without insulin) (ThermoFisher Scientific), 1% GlutaMAX (ThermoFisher Scientific), 200 µM L-ascorbic acid 2-phosphate sesquimagnesium salt hydrate and 1% Penicillin/Streptomycin (ThermoFisher Scientific), 1 µmol/L Glucose, and 100 µmol/L Palmitic acid (conjugated to bovine serum albumin within B27 supplement by incubating for 2 hours at 37ºC, Sigma) with changes every 2-3 days. For chronic doxycycline treatment experiments, hCOs were generated from CHIR/XAV cardiac-directed differentiations (as described in (Friedman et al., 2018)), treated with doxycycline from the onset of differentiation (continuing through physiological assessment), and maintained in CNTL media conditions after hCO fabrication. **Physiology Analysis:** Pole deflection was used to approximate the force of contraction as per (Mills et al., 2017a). A Leica Dmi8 inverted high content Imager was used to capture a 10 second time-lapse of each hCO contracting in real time at 37ºC. Custom batch processing files were written in MATLAB R2013a (Mathworks) to convert the stacked TIFF files to AVI, track the pole movement (using vision.PointTracker), determine the contractile parameters, produce a force-time figure, and export the batch data to an Excel (Microsoft) spreadsheet. **Whole-Mount Immunostaining:** hCOs were fixed for 60 minutes with 1% paraformaldehyde (Sigma) at 22ºC and washed 3X with PBS, after which they were incubated with Mouse anti-α-Actinin (Sigma, Cat.#A7811; RRID: AB_476766) at 1:1000 and Rabbit anti-Ki-67 (Clone D3B5, Cell Signaling Technologies, Cat.#9129; RRID: AB_2687446) at 1:400 in blocking buffer (5% FBS and 0.2% Triton X-100 [Sigma] in PBS) overnight at 4ºC. Cells were then washed in blocking buffer 2X for 2 hours and subsequently incubated with Goat anti-Mouse Alexa Fluor 555 (ThermoFisher Scientific, Cat.#A-21235; RRID: AB_2535804) and Goat anti-Rabbit Alexa Fluor 488 (ThermoFisher Scientific, Cat.#A-11008; RRID: AB_ 143165) (both at 1:400) and Hoechst 33342 (1:1000) overnight at 4ºC. They were washed in blocking buffer 2X for 2 hours and imaged *in situ* or mounted on microscope slides using Fluoromount-G (Southern Biotech). **Immunostaining Analysis:** hCOs were imaged using a Leica Dmi8 high content imaging microscope. Custom batch processing files were written in MATLAB R2013a (Mathworks) to remove the background, calculate the image intensity, and export the batch data to an Excel (Microsoft) spreadsheet. For cell cycle analysis experiments three random fields of view were imaged and manually quantified for proliferation. These were added together to calculate the percentage of hiPSC-CM cell cycle activity in each hCO.

### Flow Cytometry

As performed previously(Palpant et al., 2015), cells were fixed with 4% Paraformaldehyde (Sigma, Cat.#158127). Each sample was stained using cardiac troponin T (ThermoFisher, Cat#MA5-12960; RRID: AB_11000742) at 1:200 or a mouse IgG isotype control (BioRad, Cat.#1706516; RRID: AB_11125547) at 1:200 in 0.75% Saponin (Sigma, Cat.#S7900) for 30 minutes at 22ºC. Cells were subsequently washed with 200 µL 0.75% Saponin 2X at 22ºC. Each sample was stained in darkness with Goat anti-Mouse Alexa Fluor 488 (ThermoFisher, Cat.#A-11001; RRID:AB_2534069) at 1:200 in 0.75% Saponin for 30 minutes at 22ºC. Cells were washed with 200 µL 5% FBS in PBS 2X at 22ºC and subsequently analyzed using a BD FACSCANTO II (Becton Dickinson, San Jose, CA) with FACSDiva software (BD Biosciences). Data analysis was performed using FlowJo (Tree Star, Ashland, Oregon).

### Zebrafish Strains, Husbandry, and Transgenesis

All zebrafish experiments were performed in accordance with protocols approved by the Animal and Human Ethics Committee of The University of Queensland (UQ) (IMB/030/16/NHMRC/ARCHF). Zebrafish (*Danio rerio*) were maintained at 28°C water temperature and the light cycle was maintained at 14:10 hours (light:dark). We used the zebrafish transgenic line *Tg(cmlc2:Gal4-RFP)*(Bennett et al., 2013) (also known as *Tg(myl7:Gal4)* and *Tg(cmlc2:GAL4)*). Zebrafish work followed the guidelines of the animal ethics committee at the University of Queensland. All the embryos used in this work were generated by natural spawning and incubated in E3 medium at 28.5°C. **DNA Constructs and Transgenesis:** The *10xUAS:hopx-pA* construct was generated using PCR amplified *hopx* cDNA cloned into a middle entry (ME) vector (pDONR-221) using Gateway technology (ThermoFisher). Subsequent LR reactions were performed combining p5E-10xUAS, pME-*hopx*, and p3E-polyA into pDestTol2pAC (containing an α-crystallin promoter driving GFP in the zebrafish eye) to generate the *10xUAS:hopx-pA* DNA construct. To establish transgenic lines, these constructs (25-100 ng/µL) were separately injected with *tol2* transposase mRNA (25 ng/µL) into the cytoplasm of one-cell stage embryos. After two days, injected embryos were scored for selectable markers, raised to adulthood, and outcrossed to *Tu* zebrafish to screen for germline transmission and establish stable transgenic lines. Both male and females from these stable lines were crossed the *Tg(myl7-RFP:Gal4)* line to generate clutches of zebrafish with cardiomyocyte-specific *hopx* overexpression based on expression of cardiac RFP and eye GFP and control siblings expressing either cardiac RFP or eye GFP, but not both. **Assessment of Zebrafish Cardiac Physiology:** Assessment of zebrafish cardiac physiology was performed as previously described(Bagatto and Burggren, 2006; Hoage et al., 2012). Briefly, starting at 24 hours post fertilization, embryonic zebrafish were maintained in E3 media containing 500 µM 1-Phenyl-2-thiourea (PTU) to prevent pigment formation. At five days post fertilization zebrafish were live mounted in 1% agarose (wt/vol) and placed upon to the stage of an inverted deconvolution microscope (Nikon Ti-E) immediately prior to video recording using a monochromatic camera(Hamamatsu Flash 4.0 sCMOS). Three independent frames from each video were analyzed and averaged using FIJI (NIH) and Excel (Microsoft). **Heart Resection Model:** Heart injury experiments were performed as previously described (Poss et al., 2002). Briefly, the heart was exposed by open chest surgery under anesthesia by 0.02% Tricaine and 20% of the apical ventricle was subsequently resected using micro-scissors. Zebrafish recovered from the anesthetic in fresh water and were returned to the aquarium system. **Histology and Immunohistochemistry:** Samples were sectioned, stained, and imaged were performed as previously described (Hui et al., 2017; Sugimoto et al., 2017). Hearts harvested from *hopx* OE or control zebrafish on day 7 after the resection were fixed in 3% PFA, dehydrated in 30% sucrose, and frozen in freezing medium (TFM; Leica Biosystems, Wetzlar, Germany). Serial 10 µm sections were cut using cryostat (Leica). The sections were stained with primary antibodies against Mouse anti-PCNA (Clone PC10, Sigma, Cat.#P8825; RRID: AB_477413) and Rabbit anti-Mef2 (Clone C-21, Santa Cruz, Cat.#SC-313; RRID: AB_631920) and subsequently stained with secondaries antibodies Donkey anti-Mouse IgG conjugated Alexa Fluor 488 (ThermoFisher, Cat.# A-21202; RRID: AB_141607) and anti-Rabbit IgG conjugated Alexa Fluor 555 (ThermoFisher, Cat.# A-31572; RRID: AB_162543). **Cardiomyocyte Proliferation Analysis:** Cardiomyocyte proliferation analysis was performed as previously described(Hui et al., 2017; Sugimoto et al., 2017). All images for quantification and representation were taken using a Zeiss Axio imager M1 (Carl Zeiss AG) fluorescent microscope with a 10x and 20x objective. The numbers of PCNA^+^/Mef2^+^ cells were manually counted on the injury border zone (967 × 267 pixels) using FIJI software (NIH). The percentages of PCNA^+^/Mef2^+^ cells were determined from 3-4 sections and the percentages were averaged to obtain a cardiomyocyte proliferation percentage for each heart.

### Genome-Wide Association Studies

#### Analysis of GWAS studies

To assess whether the HOPX regulatory network associates with cardiovascular structure in healthy individuals, we performed gene-based tests on GWAS summary data using fastBAT(Bakshi et al., 2016) and SMR(Zhu et al., 2016) as implemented in the Complex-Traits Genetics Virtual Lab (CTG-VL(Cuellar-Partida et al., 2019)). We defined the HOPX regulatory network as genes which are directly bound by HOPX (3,467 genes) and/or transcriptionally dependent on HOPX (650 genes) (**Table S3**). fastBAT tests the aggregated effects of a set of genetic variants within or close to (± 50 kb) each tested gene using a set-based association approach which accounts for the correlation between genetic variants (i.e. linkage disequilibrium). This provides a more powerful approach over single-variant tests. SMR (Summary-data-based Mendelian Randomization) tests for association between the expression of a gene and a complex trait of interest in a trait-relevant tissue. All SMR analyses were performed using eQTLs identified from left ventricular heart tissue in GTEx release v7.

We performed these analyses using GWAS summary data of MRI analysis of left ventricle structural traits (N = 36,041) from the UK Biobank GWAS database(Pirruccello et al., 2020). Specifically, left ventricular end-diastolic volume, left ventricular end-diastolic volume body surface area (BSA) indexed, left ventricular ejection fraction, left ventricular end-systolic volume, left ventricular end-systolic volume BSA indexed, stroke volume and stroke volume BSA indexed. To determine whether there was enrichment of the HOPX regulatory network in the associated genes we performed a Chi-squared test or a Fisher’s exact test depending on the frequencies in the contingency table.

### Statistics

Unless otherwise indicated, unpaired t-test were performed with Welch’s correction for comparisons of two groups using GraphPad Prism 8 software. Asterisks indicate statistical significance (P<0.05). Not significant (NS) indicates P>0.05.

## REFERENCES

Alexanian, M., Maric, D., Jenkinson, S.P., Mina, M., Friedman, C.E., Ting, C.C., Micheletti, R., Plaisance, I., Nemir, M., Maison, D., et al. (2017). A transcribed enhancer dictates mesendoderm specification in pluripotency. Nat. Commun. 8, 1806.

Asp, M., Giacomello, S., Larsson, L., Wu, C., Fürth, D., Qian, X., Wärdell, E., Custodio, J., Reimegård, J., Salmén, F., et al. (2019). A Spatiotemporal Organ-Wide Gene Expression and Cell Atlas of the Developing Human Heart. Cell 179, 1647–1660.e19.

Aughey, G.N., Cheetham, S.W., and Southall, T.D. (2019). DamID as a versatile tool for understanding gene regulation. Development 146, dev173666.

Bagatto, B., and Burggren, W.W. (2006). A three-dimensional functional assessment of heart and vessel development in the larva of the zebrafish (Danio rerio). Physiol. Biochem. Zool. 79, 194–201.

Bakshi, A., Zhu, Z., Vinkhuyzen, A.A., Hill, W.D., McRae, A.F., Visscher, P.M., and Yang, J. (2016). Fast set-based association analysis using summary data from GWAS identifies novel gene loci for human complex traits. Sci. Rep. 6, 32894.

Bennett, J.S., Stroud, D.M., Becker, J.R., and Roden, D.M. (2013). Proliferation of embryonic cardiomyocytes in zebrafish requires the sodium channel scn5Lab. Genesis 51, 562–574.

Berg, D.A., Su, Y., Jimenez-Cyrus, D., Patel, A., Huang, N., Morizet, D., Lee, S., Shah, R., Ringeling, F.R., Jain, R., et al. (2019). A Common Embryonic Origin of Stem Cells Drives Developmental and Adult Neurogenesis. Cell 177, 654–668.e15.

Burridge, P.W., Matsa, E., Shukla, P., Lin, Z.C., Churko, J.M., Ebert, A.D., Lan, F., Diecke, S., Huber, B., Mordwinkin, N.M., et al. (2014). Chemically defined generation of human cardiomyocytes. Nat. Methods 11, 855–860.

Cheetham, S.W., Gruhn, W.H., van den Ameele, J., Krautz, R., Southall, T.D., Kobayashi, T., Surani, M.A., and Brand, A.H. (2018). Targeted DamID reveals differential binding of mammalian pluripotency factors. Development 145, dev170209.

Chen, F., Kook, H., Milewski, R., Gitler, A.D., Lu, M.M., Li, J., Nazarian, R., Schnepp, R., Jen, K., Biben, C., et al. (2002). Hop is an unusual homeobox gene that modulates cardiac development. Cell 110, 713–723.

Chen, Y., Yang, L., Cui, T., Pacyna-Gengelbach, M., and Petersen, I. (2015). HOPX is methylated and exerts tumour-suppressive function through Ras-induced senescence in human lung cancer. J. Pathol. 235, 397–407.

Cuellar-Partida, G., Lundberg, M., Kho, P.F., D’Urso, S., Gutierrez-Mondragon, L.F., and Hwang, L.-D. (2019). Complex-Traits Genetics Virtual Lab: A community-driven web platform for post-GWAS analyses. BioRxiv 518027.

DeLaughter, D.M., Bick, A.G., Wakimoto, H., McKean, D., Gorham, J.M., Kathiriya, I.S., Hinson, J.T., Homsy, J., Gray, J., Pu, W., et al. (2016). Single-Cell Resolution of Temporal Gene Expression during Heart Development. Dev. Cell 39, 480–490.

Eulalio, A., Mano, M., Dal Ferro, M., Zentilin, L., Sinagra, G., Zacchigna, S., and Giacca, M. (2012). Functional screening identifies miRNAs inducing cardiac regeneration. Nature 492, 376–381.

Friedman, C.E., Nguyen, Q., Lukowski, S.W., Helfer, A., Chiu, H.S., Miklas, J., Levy, S., Suo, S., Han, J.J., Osteil, P., et al. (2018). Single-Cell Transcriptomic Analysis of Cardiac Differentiation from Human PSCs Reveals HOPX-Dependent Cardiomyocyte Maturation. Cell Stem Cell 23, 586–598.e8.

Gorkin, D.U., Barozzi, I., Zhao, Y., Zhang, Y., Huang, H., Lee, A.Y., Li, B., Chiou, J., Wildberg, A., Ding, B., et al. (2020). An atlas of dynamic chromatin landscapes in mouse fetal development. Nature 583, 744–751.

Haberle, V., Forrest, A.R., Hayashizaki, Y., Carninci, P., and Lenhard, B. (2015). CAGEr: precise TSS data retrieval and high-resolution promoterome mining for integrative analyses. Nucleic Acids Res. 43, e51.

van Hengel, J., Van den Vreken, N., and Aalders, J. (2017). Maturation of human pluripotent stem cell-derived cardiomyocytes. In COST Action CA16119 CellFit: In Vitro 3-D Total Cell Guidance and Fitness,.

Hoage, T., Ding, Y., and Xu, X. (2012). Quantifying cardiac functions in embryonic and adult zebrafish. Methods Mol. Biol. 843, 11–20.

Huang, da W., Sherman, B.T., and Lempicki, R.A. (2009). Systematic and integrative analysis of large gene lists using DAVID bioinformatics resources. Nat. Protoc. 4, 44–57.

Hudson, J.E., and Zimmermann, W.H. (2011). Tuning Wnt-signaling to enhance cardiomyogenesis in human embryonic and induced pluripotent stem cells. J. Mol. Cell. Cardiol. 51, 277–279.

Hudson, J., Titmarsh, D., Hidalgo, A., Wolvetang, E., and Cooper-White, J. (2012). Primitive cardiac cells from human embryonic stem cells. Stem Cells Dev. 21, 1513–1523.

Hui, S.P., Sheng, D.Z., Sugimoto, K., Gonzalez-Rajal, A., Nakagawa, S., Hesselson, D., and Kikuchi, K. (2017). Zebrafish Regulatory T Cells Mediate Organ-Specific Regenerative Programs. Dev. Cell 43, 659–672.e5.

Jain, R., Li, D., Gupta, M., Manderfield, L.J., Ifkovits, J.L., Wang, Q., Liu, F., Liu, Y., Poleshko, A., Padmanabhan, A., et al. (2015). HEART DEVELOPMENT. Integration of Bmp and Wnt signaling by Hopx specifies commitment of cardiomyoblasts. Science 348, aaa6071.

Jopling, C., Sleep, E., Raya, M., Martí, M., Raya, A., and Izpisúa Belmonte, J.C. (2010). Zebrafish heart regeneration occurs by cardiomyocyte dedifferentiation and proliferation. Nature 464, 606–609.

Karbassi, E., Fenix, A., Marchiano, S., Muraoka, N., Nakamura, K., Yang, X., and Murry, C.E. (2020). Cardiomyocyte maturation: advances in knowledge and implications for regenerative medicine. Nat. Rev. Cardiol. 2020/02/03.

Kikuchi, K., Holdway, J.E., Werdich, A.A., Anderson, R.M., Fang, Y., Egnaczyk, G.F., Evans, T., Macrae, C.A., Stainier, D.Y., and Poss, K.D. (2010). Primary contribution to zebrafish heart regeneration by gata4(+) cardiomyocytes. Nature 464, 601–605.

King, E.A., Davis, J.W., and Degner, J.F. (2019). Are drug targets with genetic support twice as likely to be approved? Revised estimates of the impact of genetic support for drug mechanisms on the probability of drug approval. PLoS Genet. 15, e1008489.

Kook, H., Lepore, J.J., Gitler, A.D., Lu, M.M., Yung, W.W.-M., Mackay, J., Zhou, R., Ferrari, V., Gruber, P., and Epstein, J.A. (2003). Cardiac hypertrophy and histone deacetylase-dependent transcriptional repression mediated by the atypical homeodomain protein Hop. J. Clin. Invest. 112.

Lescroart, F., Wang, X., Lin, X., Swedlund, B., Gargouri, S., Sànchez-Dànes, A., Moignard, V., Dubois, C., Paulissen, C., Kinston, S., et al. (2018). Defining the earliest step of cardiovascular lineage segregation by single-cell RNA-seq. Science 2018/01/25.

Li, G., Xu, A., Sim, S., Priest, J.R., Tian, X., Khan, T., Quertermous, T., Zhou, B., Tsao, P.S., Quake, S.R., et al. (2016). Transcriptomic Profiling Maps Anatomically Patterned Subpopulations among Single Embryonic Cardiac Cells. Dev. Cell 39, 491–507.

Lian, X., Hsiao, C., Wilson, G., Zhu, K., Hazeltine, L.B., Azarin, S.M., Raval, K.K., Zhang, J., Kamp, T.J., and Palecek, S.P. (2012). Robust cardiomyocyte differentiation from human pluripotent stem cells via temporal modulation of canonical Wnt signaling. Proc. Natl. Acad. Sci. U. S. A. 109, E1848–E1857.

Lin, C.-C., Yao, C.-Y., Hsu, Y.-C., Hou, H.-A., Yuan, C.-T., Li, Y.-H., Kao, C.-J., Chuang, P.- H., Chiu, Y.-C., Chen, Y., et al. (2020). Knock-out of Hopx disrupts stemness and quiescence of hematopoietic stem cells in mice. Oncogene 39, 5112–5123.

Liu, Y., and Zhang, W. (2020). The role of HOPX in normal tissues and tumor progression. Biosci. Rep. 40.

Loh, K.M., Chen, A., Koh, P.W., Deng, T.Z., Sinha, R., Tsai, J.M., Barkal, A.A., Shen, K.Y., Jain, R., Morganti, R.M., et al. (2016). Mapping the Pairwise Choices Leading from Pluripotency to Human Bone, Heart, and Other Mesoderm Cell Types. Cell 166, 451–467.

Love, M.I., Huber, W., and Anders, S. (2014). Moderated estimation of fold change and dispersion for RNA-seq data with DESeq2. Genome Biol. 15, 550.

Luna-Zurita, L., Stirnimann, C.U., Glatt, S., Kaynak, B.L., Thomas, S., Baudin, F., Samee, M.A.H., He, D., Small, E.M., Mileikovsky, M., et al. (2016). Complex Interdependence Regulates Heterotypic Transcription Factor Distribution and Coordinates Cardiogenesis. Cell 164, 999–1014.

Machanick, P., and Bailey, T.L. (2011). MEME-ChIP: motif analysis of large DNA datasets. Bioinformatics 27, 1696–1697.

Mandegar, M.A., Huebsch, N., Frolov, E.B., Shin, E., Truong, A., Olvera, M.P., Chan, A.H., Miyaoka, Y., Holmes, K., Spencer, C.I., et al. (2016). CRISPR Interference Efficiently Induces Specific and Reversible Gene Silencing in Human iPSCs. Cell Stem Cell 18, 541– 553.

Marshall, O.J., and Brand, A.H. (2015). Damidseq-pipeline: An automated pipeline for processing DamID sequencing datasets. Bioinformatics 31, 3371–3373.

Marshall, O.J., Southall, T.D., Cheetham, S.W., and Brand, A.H. (2016). Cell-type-specific profiling of protein-DNA interactions without cell isolation using targeted DamID with next- generation sequencing. Nat. Protoc. 11, 1586–1598.

McLean, C.Y., Bristor, D., Hiller, M., Clarke, S.L., Schaar, B.T., Lowe, C.B., Wenger, A.M., and Bejerano, G. (2010). GREAT improves functional interpretation of cis-regulatory regions. Nat. Biotechnol. 28, 495–501.

Mills, R.J., Titmarsh, D.M., Koenig, X., Parker, B.L., Ryall, J.G., Quaife-Ryan, G.A., Voges, H.K., Hodson, M.P., Ferguson, C., Drowley, L., et al. (2017a). Functional screening in human cardiac organoids reveals a metabolic mechanism for cardiomyocyte cell cycle arrest. Proc. Natl. Acad. Sci. U. S. A. 114, E8372–E8381.

Mills, R.J., Voges, H.K., Porrello, E.R., and Hudson, J.E. (2017b). Cryoinjury Model for Tissue Injury and Repair in Bioengineered Human Striated Muscle. Methods Mol. Biol. 1668, 209–224.

Mills, R.J., Parker, B.L., Quaife-Ryan, G.A., Voges, H.K., Needham, E.J., Bornot, A., Ding, M., Andersson, H., Polla, M., Elliott, D.A., et al. (2019). Drug Screening in Human PSC-Cardiac Organoids Identifies Pro-proliferative Compounds Acting via the Mevalonate Pathway. Cell Stem Cell 24, 895–907 e6.

Nicin, L., Abplanalp, W.T., Schänzer, A., Sprengel, A., John, D., Mellentin, H., Tombor, L., Keuper, M., Ullrich, E., Klingel, K., et al. (2021). Single Nuclei Sequencing Reveals Novel Insights into the Regulation of Cellular Signatures in Children with Dilated Cardiomyopathy. Circulation.

Ock, S., Lee, W.S., Ahn, J., Kim, H.M., Kang, H., Kim, H.-S., Jo, D., Abel, E.D., Lee, T.J., and Kim, J. (2016). Deletion of IGF-1 Receptors in Cardiomyocytes Attenuates Cardiac Aging in Male Mice. Endocrinology 157, 336–345.

Palpant, N.J., Pabon, L., Rabinowitz, J.S., Hadland, B.K., Stoick-Cooper, C.L., Paige, S.L., Bernstein, I.D., Moon, R.T., and Murry, C.E. (2013). Transmembrane protein 88: a Wnt regulatory protein that specifies cardiomyocyte development. Development 140, 3799–3808.

Palpant, N.J., Pabon, L., Roberts, M., Hadland, B., Jones, D., Jones, C., Moon, R.T., Ruzzo, W.L., Bernstein, I., Zheng, Y., et al. (2015). Inhibition of beta-catenin signaling respecifies anterior-like endothelium into beating human cardiomyocytes. Development 142, 3198– 3209.

Palpant, N.J., Wang, Y., Hadland, B., Zaunbrecher, R.J., Redd, M., Jones, D., Pabon, L., Jain, R., Epstein, J., Ruzzo, W.L., et al. (2017a). Chromatin and Transcriptional Analysis of Mesoderm Progenitor Cells Identifies HOPX as a Regulator of Primitive Hematopoiesis. Cell Rep. 20, 1597–1608.

Palpant, N.J., Pabon, L., Friedman, C.E., Roberts, M., Hadland, B., Zaunbrecher, R.J., Bernstein, I., Zheng, Y., and Murry, C.E. (2017b). Generating high-purity cardiac and endothelial derivatives from patterned mesoderm using human pluripotent stem cells. Nat. Protoc. 12, 15–31.

Pavlova, O., Lefort, K., Mariotto, A., Huber, M., and Hohl, D. (2021). Homeodomain only protein X (HOPX) has an oncogenic activity during squamous skin carcinogenesis. J. Invest. Dermatol.

Pirruccello, J.P., Bick, A., Wang, M., Chaffin, M., Friedman, S., Yao, J., Guo, X., Venkatesh, B.A., Taylor, K.D., Post, W.S., et al. (2020). Analysis of cardiac magnetic resonance imaging in 36,000 individuals yields genetic insights into dilated cardiomyopathy. Nat. Commun. 11, 2254.

Poss, K.D., Wilson, L.G., and Keating, M.T. (2002). Heart regeneration in zebrafish. Science 298, 2188–2190.

Quinlan, A.R., and Hall, I.M. (2010). BEDTools: a flexible suite of utilities for comparing genomic features. Bioinformatics 26, 841–842.

Ramírez, F., Dündar, F., Diehl, S., Grüning, B.A., and Manke, T. (2014). deepTools: a flexible platform for exploring deep-sequencing data. Nucleic Acids Res. 42, W187–W191.

Redd, M.A., Scheuer, S.E., Saez, N.J., Yoshikawa, Y., Chiu, H.S., Gao, L., Hicks, M., Villanueva, J.E., Joshi, Y., Chow, C.Y., et al. (2021). Therapeutic Inhibition of Acid-Sensing Ion Channel 1a Recovers Heart Function After Ischemia–Reperfusion Injury. Circulation 144, 947–960.

Sala, L., van Meer, B.J., Tertoolen, L.G.J., Bakkers, J., Bellin, M., Davis, R.P., Denning, C., Dieben, M.A.E., Eschenhagen, T., Giacomelli, E., et al. (2018). MUSCLEMOTION: A Versatile Open Software Tool to Quantify Cardiomyocyte and Cardiac Muscle Contraction In Vitro and In Vivo. Circ. Res. 122, e5–e16.

Shim, W.J., Sinniah, E., Xu, J., Vitrinel, B., Alexanian, M., Andreoletti, G., Shen, S., Sun, Y., Balderson, B., Boix, C., et al. (2020). Conserved Epigenetic Regulatory Logic Infers Genes Governing Cell Identity. Cell Syst 11, 625–639 e13.

Shin, C.H., Liu, Z.P., Passier, R., Zhang, C.L., Wang, D.Z., Harris, T.M., Yamagishi, H., Richardson, J.A., Childs, G., and Olson, E.N. (2002). Modulation of cardiac growth and development by HOP, an unusual homeodomain protein. Cell 110, 725–735.

Sim, C.B., Phipson, B., Ziemann, M., Rafehi, H., Mills, R.J., Watt, K.I., Abu-Bonsrah, K.D., Kalathur, R.K.R., Voges, H.K., Dinh, D.T., et al. (2021). Sex-Specific Control of Human Heart Maturation by the Progesterone Receptor. Circulation 143, 1614–1628.

van Steensel, B., and Henikoff, S. (2000). Identification of in vivo DNA targets of chromatin proteins using tethered dam methyltransferase. Nat. Biotechnol. 18, 424–428.

Stempor, P., and Ahringer, J. (2016). SeqPlots - Interactive software for exploratory data analyses, pattern discovery and visualization in genomics. Wellcome Open Research 1, 14.

Stuart, T., Butler, A., Hoffman, P., Hafemeister, C., Papalexi, E., Mauck, W.M., 3rd, Hao, Y., Stoeckius, M., Smibert, P., and Satija, R. (2019). Comprehensive Integration of Single-Cell Data. Cell 177, 1888–1902.e21.

Sugimoto, K., Hui, S.P., Sheng, D.Z., and Kikuchi, K. (2017). Dissection of zebrafish shha function using site-specific targeting with a Cre-dependent genetic switch. Elife 6.

Tiburcy, M., Hudson, J.E., Balfanz, P., Schlick, S., Meyer, T., Chang Liao, M.L., Levent, E., Raad, F., Zeidler, S., Wingender, E., et al. (2017). Defined Engineered Human Myocardium With Advanced Maturation for Applications in Heart Failure Modeling and Repair. Circulation 135, 1832–1847.

Trivedi, C.M., Zhu, W., Wang, Q., Jia, C., Kee, H.J., Li, L., Hannenhalli, S., and Epstein, J.A. (2010). Hopx and Hdac2 interact to modulate Gata4 acetylation and embryonic cardiac myocyte proliferation. Dev. Cell 19, 450–459.

Trivedi, C.M., Cappola, T.P., Margulies, K.B., and Epstein, J.A. (2011). Homeodomain only protein x is down-regulated in human heart failure. J. Mol. Cell. Cardiol. 50, 1056–1058.

Ueno, S., Weidinger, G., Osugi, T., Kohn, A.D., Golob, J.L., Pabon, L., Reinecke, H., Moon, R.T., and Murry, C.E. (2007). Biphasic role for Wnt/beta-catenin signaling in cardiac specification in zebrafish and embryonic stem cells. Proc. Natl. Acad. Sci. U. S. A. 104, 9685–9690.

Vivien, C.J., Hudson, J.E., and Porrello, E.R. (2016). Evolution, comparative biology and ontogeny of vertebrate heart regeneration. NPJ Regen Med 1, 16012.

Voges, H.K., Mills, R.J., Elliott, D.A., Parton, R.G., Porrello, E.R., and Hudson, J.E. (2017). Development of a human cardiac organoid injury model reveals innate regenerative potential. Development 144, 1118–1127.

Yokota, T., Li, J., Huang, J., Xiong, Z., Zhang, Q., Chan, T., Ding, Y., Rau, C., Sung, K., Ren, S., et al. (2020). p38 Mitogen-activated protein kinase regulates chamber-specific perinatal growth in heart. J. Clin. Invest. 130, 5287–5301.

Zhu, Z., Zhang, F., Hu, H., Bakshi, A., Robinson, M.R., Powell, J.E., Montgomery, G.W., Goddard, M.E., Wray, N.R., Visscher, P.M., et al. (2016). Integration of summary data from GWAS and eQTL studies predicts complex trait gene targets. Nat. Genet. 48, 481–487.

