## Supplemental Information for "HOPX governs a molecular and physiological switch between cardiomyocyte progenitor and maturation gene programs"

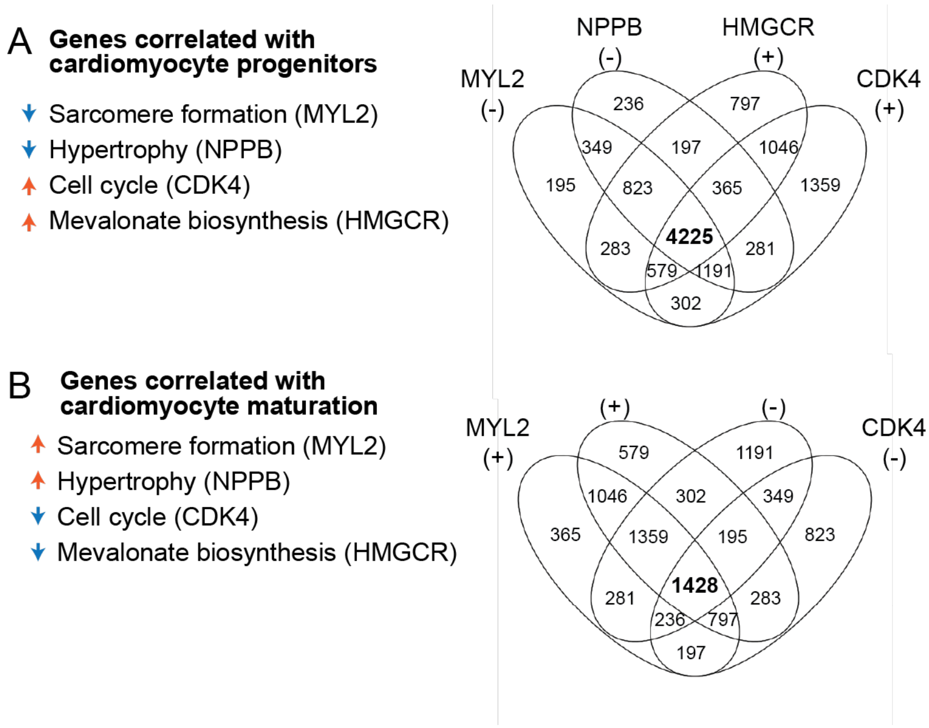


**Figure S1. Correlation analysis identifies gene programs associated with cardiac progenitor and maturation gene programs.**

(**A-B**) Analysis of gene correlations in 2,337 annotated cardiomyocytes between E6.5-P21. (**A**) Data reveal 4,225 genes negatively correlated with expression levels of *Myl2* and *Nppb* but positively correlated with expression levels of *Cdk4* and *Hmgcr.*
(**B**) Data reveal 1,428 genes positively correlated with expression levels of *Myl2* and *Nppb* but negatively correlated with expression levels of *Cdk4* and *Hmgcr.*

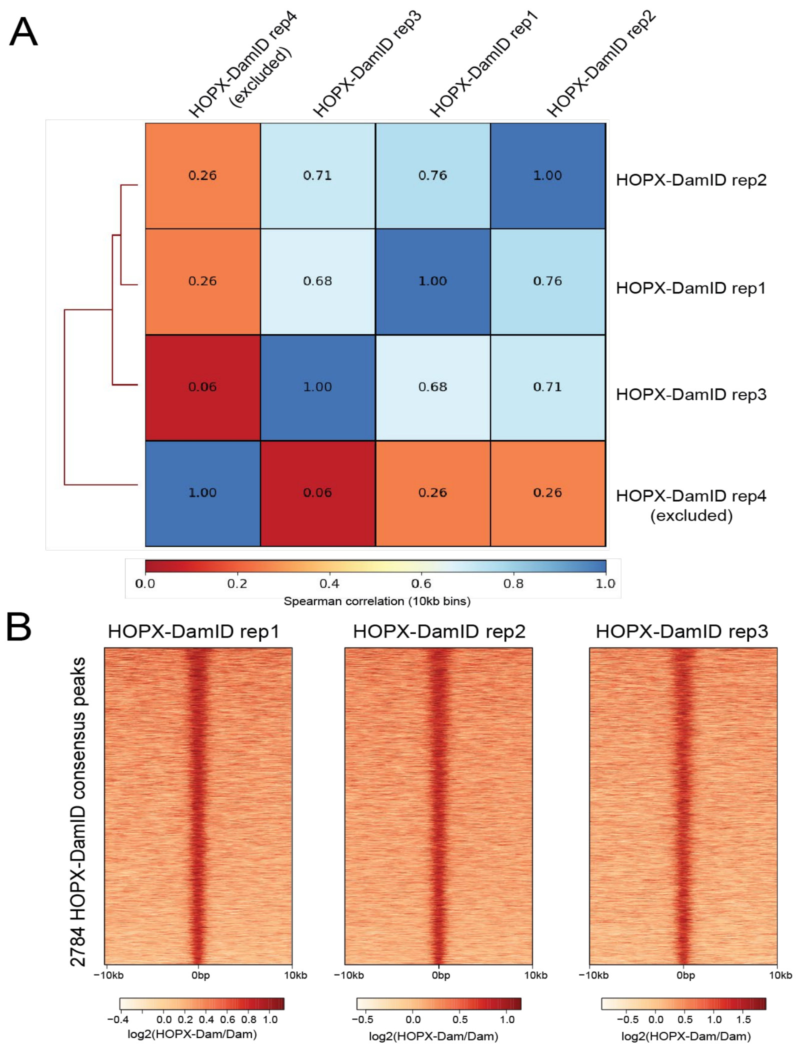


**Figure S2. DamID-sequencing quality control in hiPSC-derived cardiomyocytes.**

(**A**) Correlation of HOPX-Dam replicates over 1kb bins of the human genome. Replicate 4 is a clear outlier and was excluded. Heatmap generated with deepTools (Ramírez et al., 2014). (**B**) Heatmap demonstrating the reproducibility of HOPX-Dam signal over the consensus peaks. Heatmap generated with SeqPlots (Stempor and Ahringer, 2016).

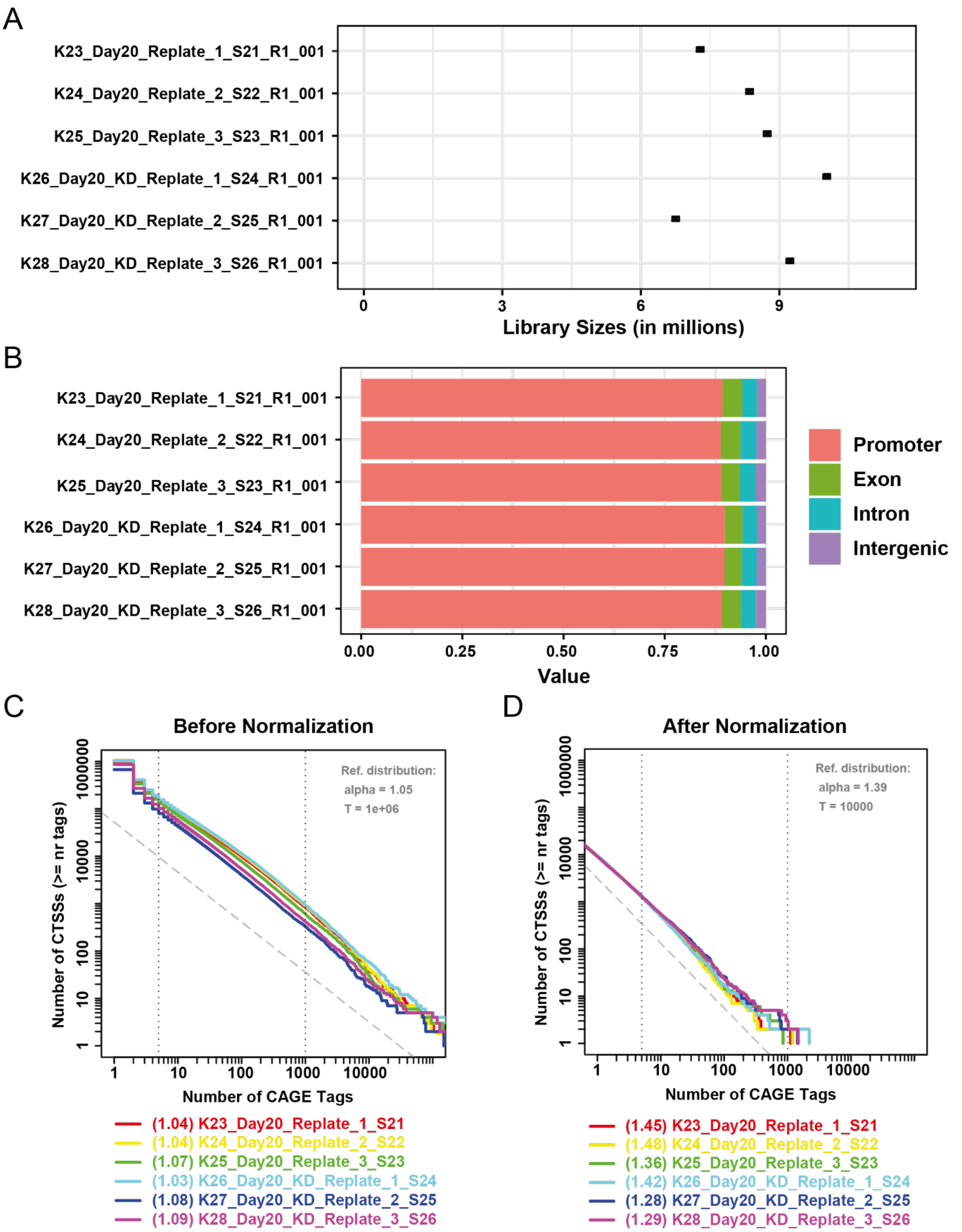


**Figure S3. Quality control of CAGE-sequencing data from hiPSC-derived cardiomyocytes.**

(**A**-**D**) Three independent biological replicates of iPSC-derived cardiomyocytes replated at day 15 and assayed at day 20 under control (K23-25) versus *HOPX* knockdown conditions (KD)

(K26-28) and analyzed for library size as total CAGE sequencing reads per sample (**A**), proportions of reads confidently mapped to the four annotated genomic features (**B**-**D**), and reverse cumulative distribution of the number of CAGE transcription start sites (CTSSs) relative to the number of CAGE sequencing reads (CAGE tags) that were mapped to the CTSSs. Y-axis shows the count of CTSSs with more than or equal to the number of corresponding CAGE tags on the X-axis. A CTSS with a higher tag count suggests the higher confidence in CTSS calling and the higher expression of the CTSS. The distribution follows a power law, which shows a reverse trend (in log 10 scale) when there is an increase in the number of CAGE tags and a decrease in the number of CTSS counts.

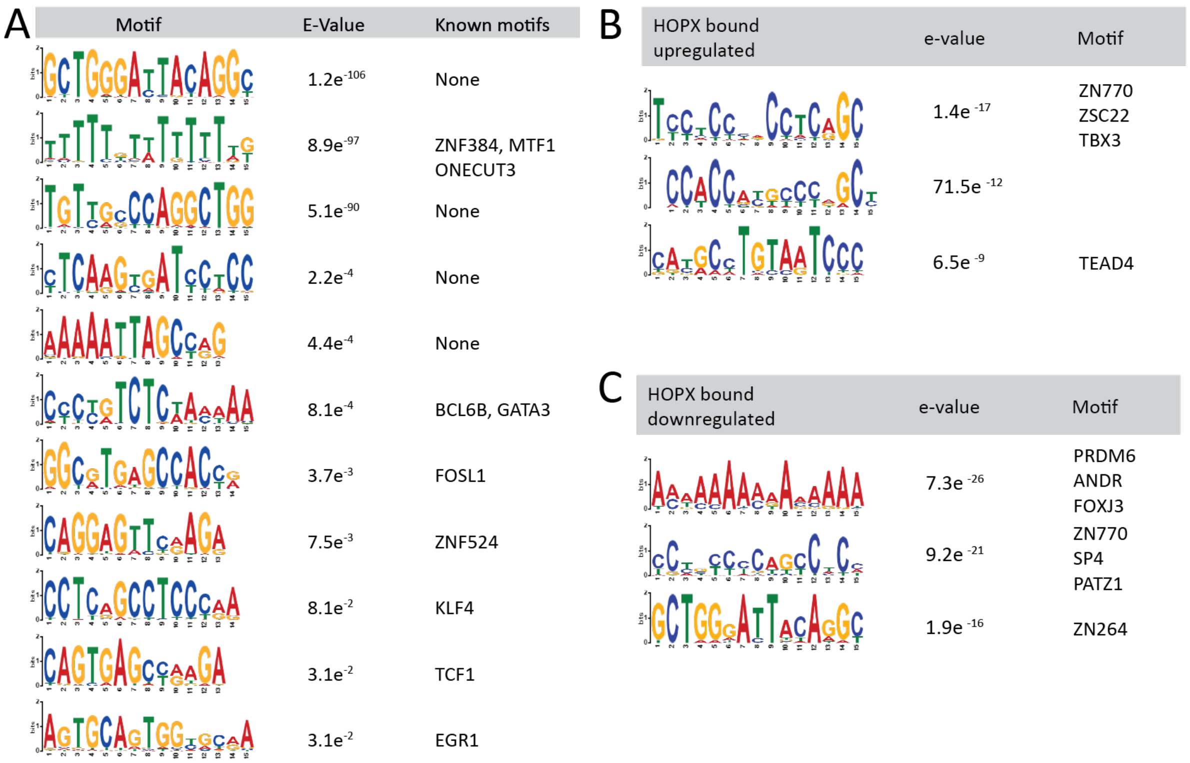


**Figure S4. Transcription factor binding site motif analysis from HOPX-Dam peaks.**

MEME-ChIP motif enrichment of all HOPX-Dam consensus peaks (**A**) and peaks associated genes that are upregulated (**B**) or downregulated (**C**) upon *HOPX* knockdown.

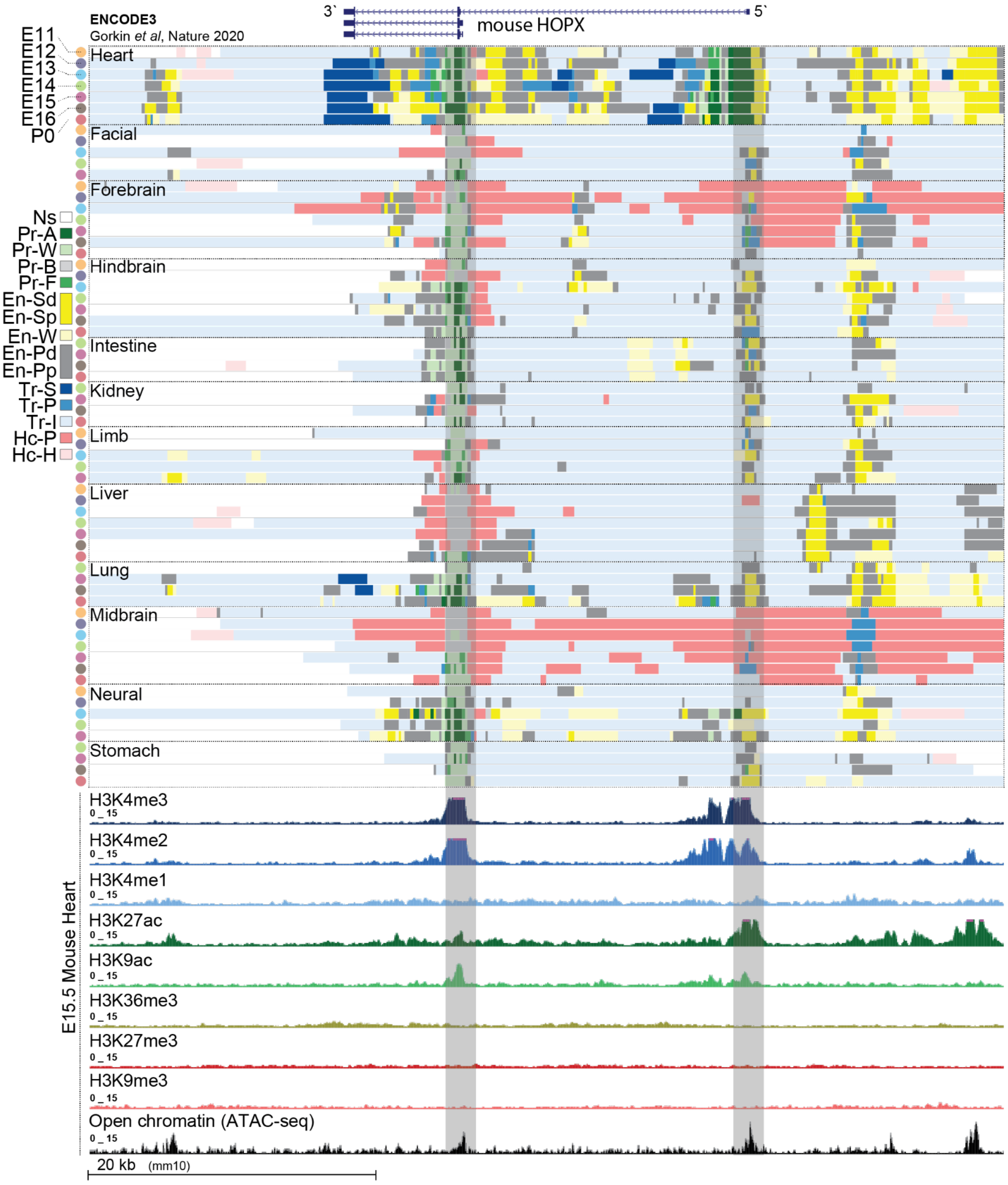


**Figure S5. ENCODE3 analysis of *HOPX* locus across diverse tissues during a time course of mouse development.**

Data display regions with no significant signal (Ns), H3K9me3-associated heterochromatin (Hc-H), polycomb-associated heterochromatin (Hc-P), transcription initiation (Tr-I), transcription permissive (Tr-P), poised TSS-proximal enhancer (En-Pp), poised TSs-distal enhancer (En-Pd), weak enhancer (En-W), strong TSS-proximal enhancer (En-Sp), strong TSS-distal enhancer (En-Sd), promoter flanking region (Pr-F), promoter bivalent (Pr-B), promoter weak (Pr-W), and promoter active (Pr-A). Raw ATAC-seq and histone-modification data tracks for heart tissue at E15.5 are shown at the bottom.

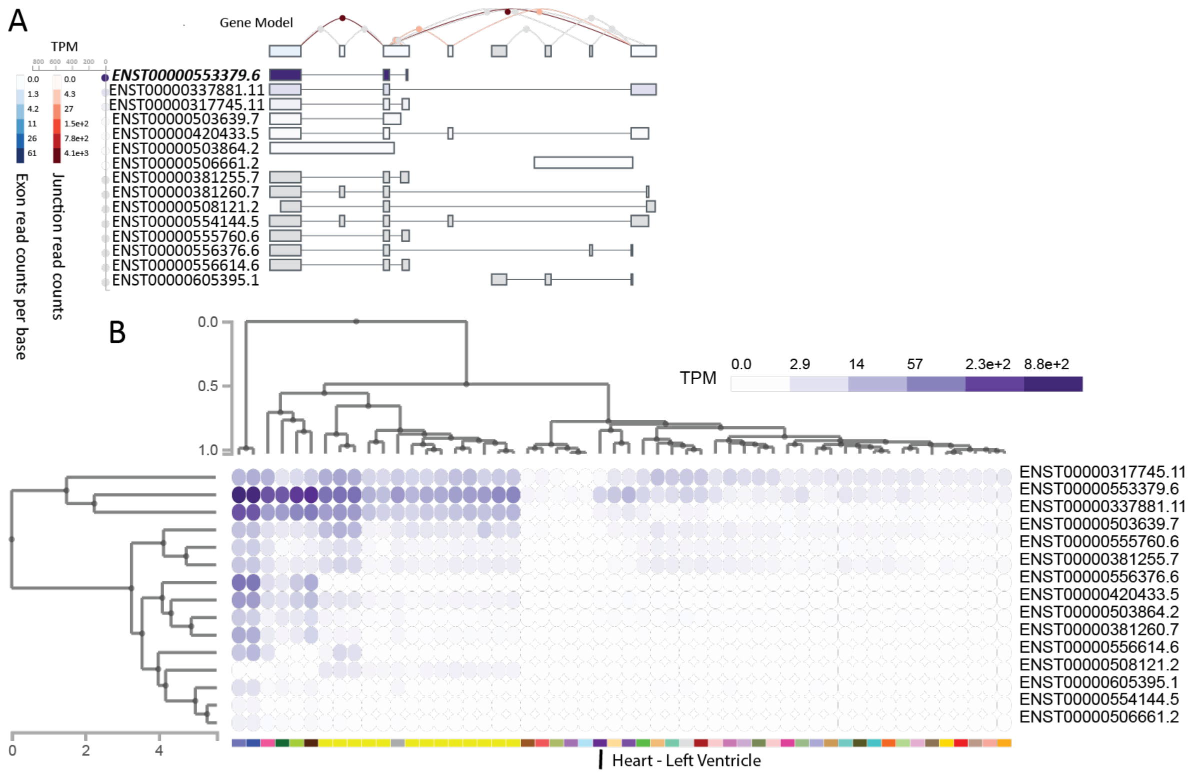


**Figure S6. GTEx analysis of *HOPX* isoforms.**

Exon expression (**A**) and isoform expression (**B**) of *HOPX*: ENSG00000171476.21 HOP homeobox [Source:HGNC Symbol;Acc:HGNC:24961].

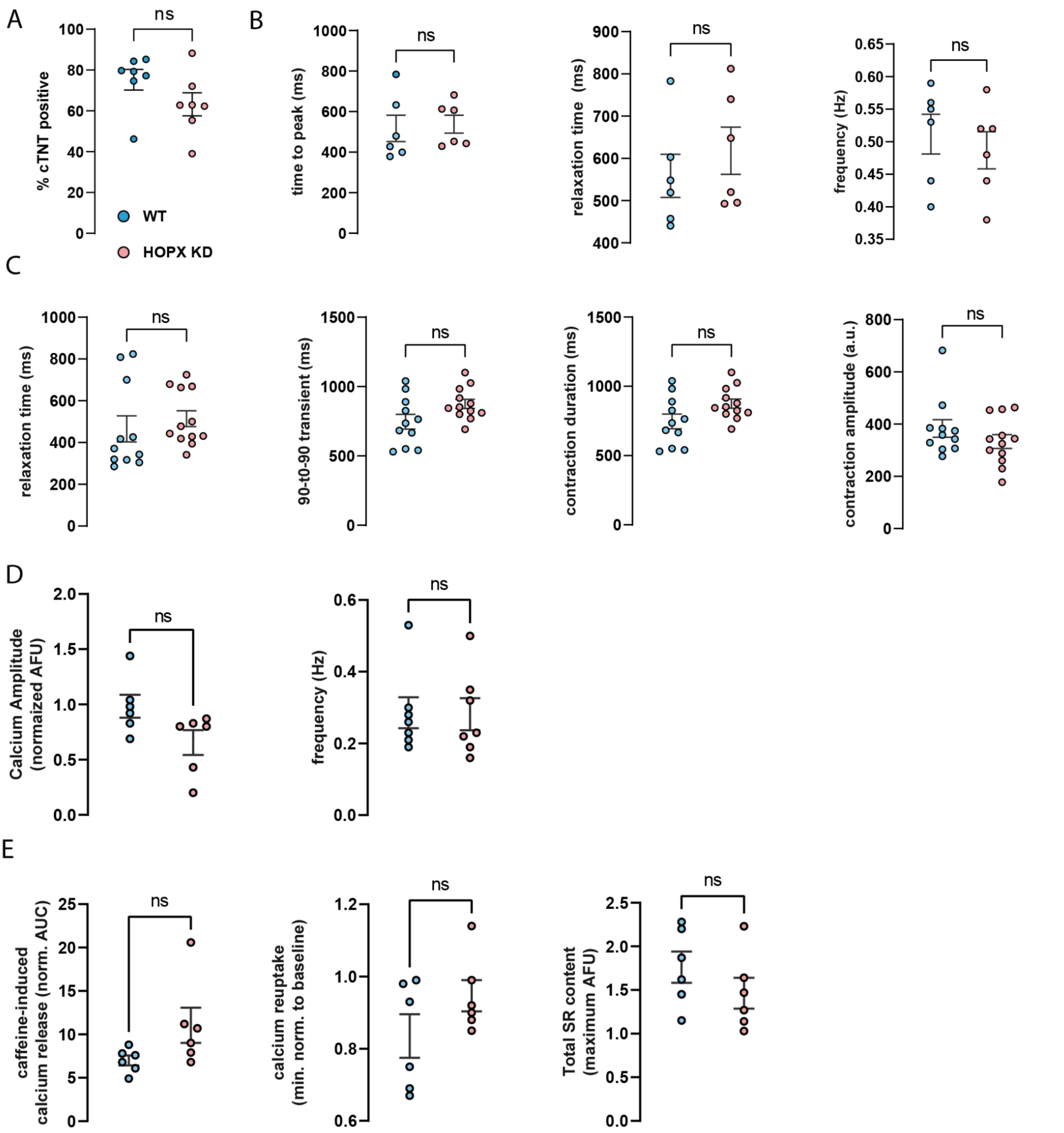


**Figure S7. iPSC-derived cardiomyocyte physiological analysis.**

(**A**) Analysis of cardiomyocyte purity in *HOPX* KD and control iPSC differentiation measured by percent cardiac troponin T by fluorescence activated cell sorting

(**B-C**) iPSC-derived cardiomyocyte contractility kinetics measured by MUSCLEMOTION at baseline (**B**) and after exposure to 10 µM noradrenaline (**C**).

(**D-E**) Calcium kinetics in monolayer derived iPSC-cardiomyocytes measured by FLIPR at baseline (**D**) and after exposure to 20 mM caffeine (**E**).

ns: not significantly different by t-test

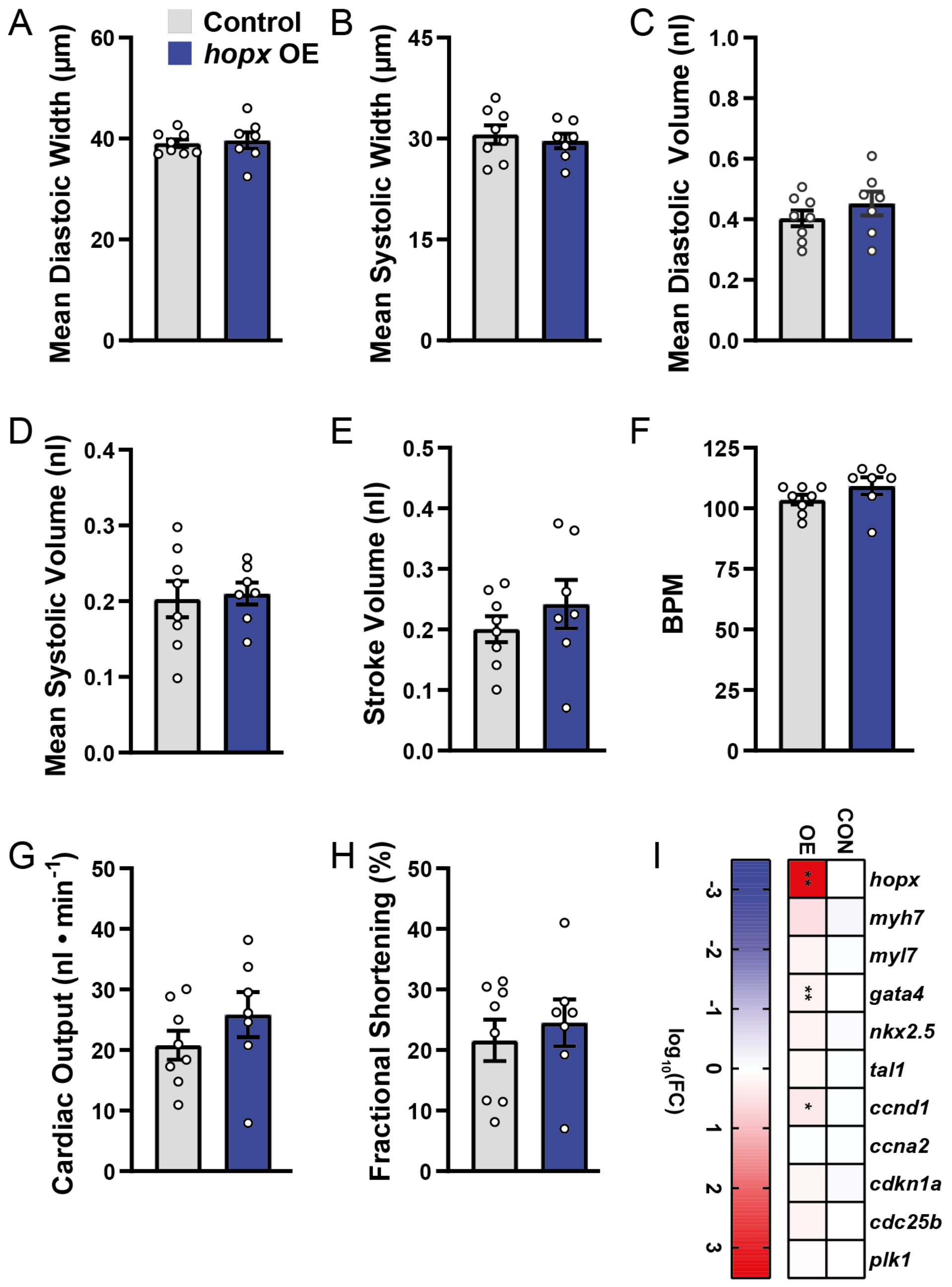


**Figure S8. Analysis of *hopx* overexpressing zebrafish.**

(**A-H**) Functional and morphometric measurements taken at 5 days post fertilization in control versus *hopx* OE including mean diastolic width (**A**), mean systolic width (**B**), mean diastolic volume (**C**), mean systolic volume (**D**), stroke volume (**E**), beats per minute (BPM) (**F**), cardiac output (**G**), and fractional shortening (**H**). ** P < 0.05 by One-Way ANOVA.

(**I**) Gene expression of candidate cardiac genes measured by quantitative RT-PCR in control versus *hopx* OE zebrafish hearts at approximately six months post fertilization. ** $P < 0.05$ by *t* test.

A **Mouse Phenotype Single KO**

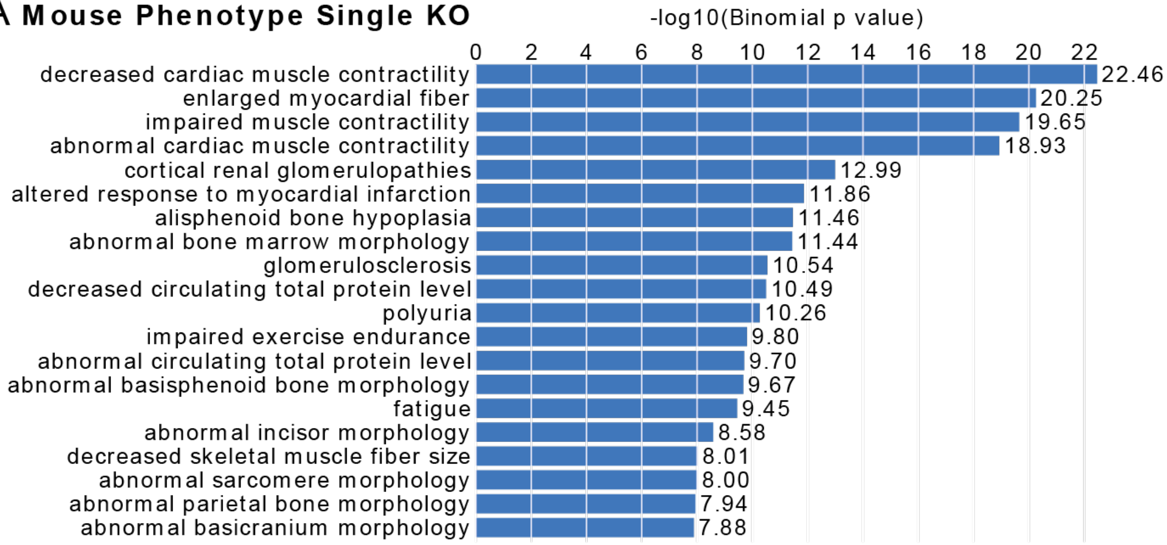


B **HOPX locus TWAS analysis**

| BEST.GWAS.ID | BEST.GWAS.Z | EQTL.ID | EQTL.GWAS.Z | TWAS.Z | TWAS.P | TRAIT |
|---|---|---|---|---|---|---|
| rs41495551 | -3.26 | rs6829867 | -0.722 | -0.58808 | 0.556 | RA maximum area |
| rs7656012 | 2.92 | rs6829867 | -0.69 | -0.80891 | 0.41857 | RA minimum area |
| rs1277272 | -3.29 | rs6829867 | 0.075 | 0.63024 | 0.52854 | RA ejection Fraction |
| rs7684253 | 5.13 | rs9968381 | 1.103 | 1.02953 | 0.303 | RV end diastolic volume |
| rs1714020 | -2.43 | rs9968381 | -0.539 | -0.12545 | 0.90017 | RV ejection fraction |
| rs7684253 | 3.09 | rs9968381 | 0.824 | 0.48061 | 0.6308 | RV end systolic volume |
| rs7684253 | 5.25 | rs9968381 | 0.755 | 0.89832 | 0.369 | RV stroke volume |
| rs2412772 | -3.35 | rs1105574 | -0.319 | 1.12851 | 0.259 | Pulmonary artery diameter |
| rs9968381 | 2.78 | rs1105574 | 1.977 | 1.723276 | 0.0848 | Pulmonary artery root diamenter |

**Figure S9. Analysis of *HOPX* in genome-wide association analysis and gene knockout phenotype databases.**

(**A**) Mouse knockout phenotypes associated with HOPX-DamID bound genes. Enrichment determined by GREAT (McLean et al., 2010).

(**B**) Analysis of *HOPX*-associated eQTLs in human GWAS data linked to cardiac structure and function parameters by MRI. Data reveal no significant associations of any *HOPX* eQTLs to volume, function, or other cardiac morphometric measurements.

**Supplemental Tables**

**Table S1. Correlation Analyses, Related to Figure 1.**

| | |
|---|---|
| Sheet 2 | GO analysis of maturation-associated genes |
| Sheet 3 | RTS analysis of maturation-associated genes |
| Sheet 4 | GO analysis of progenitor-associated genes |
| Sheet 5 | RTS analysis of progenitor-associated genes |
| Sheet 6 | *Cdk4* Correlation Analysis Using Integrated Data from Lescroart et al. 2018, Li et al. 2016, and DeLaughter et al. 2016 |
| Sheet 7 | *Hmgcr* Correlation Analysis Using Integrated Data from Lescroart et al. 2018, Li et al. 2016, and DeLaughter et al. 2016 |
| Sheet 8 | *Nppb* Correlation Analysis Using Integrated Data from Lescroart et al. 2018, Li et al. 2016, and DeLaughter et al. 2016 |
| Sheet 9 | *Myl2* Correlation Analysis Using Integrated Data from Lescroart et al. 2018, Li et al. 2016, and DeLaughter et al. 2016 |
| Sheet 10 | *Hopx* Correlation Analysis Using Integrated Data from Lescroart et al. 2018, Li et al. 2016, and DeLaughter et al. 2016 |
| Sheet 11 | *HOPX* Correlation Analysis Using Integrated Data from Asp et al. 2019 and Sim et al. 2021 |
| Sheet 12 | *HOPX* Correlation Analysis Using Data from Nicin et al. 2021 |

**Table S2. DamID-Sequencing Analyses, Related to Figures 1-3.**

| | |
|---|---|
| Sheet 2 | List of HOPX-Dam Consensus Peaks |
| Sheet 3 | GREAT Analyses of HOPX Associated Peaks |
| Sheet 4 | Gene Ontology Analyses of HOPX Peak-Associated Genes - Biological Process |
| Sheet 5 | Gene Ontology Analyses of HOPX Peak-Associated Genes - Cellular Function |
| Sheet 6 | Gene Ontology Analyses of HOPX Peak-Associated Genes - Molecular Function |
| Sheet 7 | Progenitor genes bound by HOPX |
| Sheet 8 | Gene Ontology Analysis of HOPX-Bound Progenitor-Associated Genes |

| | |
|---|---|
| Sheet 9 | Upregulated HOPX-bound progenitor genes |
| Sheet 10 | GO of upregulated HOPX-bound progenitor genes |
| Sheet 11 | Downregulated HOPX-bound progenitor genes |
| Sheet 12 | GO of downregulated HOPX-bound progenitor genes |
| Sheet 13 | Upregulated HOPX-bound maturation genes |
| Sheet 14 | GO of upregulated HOPX-bound maturation genes |
| Sheet 15 | Downregulated HOPX-bound maturation genes |
| Sheet 16 | GO of downregulated HOPX-bound maturation genes |
| Sheet 17 | Maturation genes bound by HOPX |
| Sheet 18 | Gene Ontology Analysis of HOPX-Bound Maturation-Associated Genes |

**Table S3. CAGE-sequencing Analyses, and DEP, Related to Figure 3.**

| | |
|---|---|
| Sheet 2 | TPM Matrix |
| Sheet 3 | DEG |
| Sheet 4 | GO Analyses |
| Sheet 5 | DEP |
| Sheet 6 | HOPX-Bound and Upregulated Genes |
| Sheet 7 | HOPX-Bound and Downregulated Genes |
| Sheet 8 | HOPX-Bound and Upregulated Genes GO Analysis |
| Sheet 9 | HOPX-Bound and Downregulated Genes GO Analysis |
